# Structural mechanism of mRNA decoding by mammalian GTPase GTPBP1

**DOI:** 10.1101/2025.09.26.678779

**Authors:** Denis Susorov, Anna Miścicka, Dmitrij Golovenko, Anna B. Loveland, Alexandra Zinoviev, Tatyana V. Pestova, Andrei A. Korostelev

## Abstract

GTP-binding protein 1 (GTPBP1) is a widespread translational GTPase closely related to elongation factor eEF1A. The loss of GTPBP1 leads to errors in neuronal development in animals and is associated with neurodegenerative disorders in humans. Although linked to translation and quality control mechanisms, GTPBP1 functions remain largely obscure. Similarly to eEF1A, GTPBP1 delivers cognate aminoacyl-tRNA to the ribosomal A site in a GTP-dependent manner, but GTP hydrolysis is not followed by rapid peptide bond formation, and GTPBP1-mediated elongation is slow. To establish the basis for GTPBP1 function, we determined cryo-EM structures of 80S ribosomal complexes bound to GTPBP1•aa-tRNA with GTP or the non-hydrolysable analog GDPCP. They revealed that the unique GTPBP1 architecture, including the additional eIF1/IF3-like N-terminal domain and the shoulder-interacting H-loop in place of the α2 helix of canonical GTPases, is responsible for establishing GTPBP1-specific interactions with tRNA and the ribosome, leading to slow GTPBP1 dissociation after GTP hydrolysis and thus delayed tRNA accommodation. Slow dissociation correlates with an extended proofreading stage resulting in higher accuracy of GTPBP1-mediated decoding and potentially allows GTPBP1 to elicit its putative quality control functions. GTPBP1 visualization provides the foundation for mapping and elucidating GTPBP1 mutations associated with human diseases.

## Introduction

GTP-binding protein 1 (GTPBP1) belongs to the widespread GTPBP1/GTPBP2 group of translational GTPases that are closely related to elongation factor eEF1A (and its prokaryotic homolog EF-Tu) delivering aminoacyl-tRNAs to the ribosome, eukaryotic release factor eRF3 and surveillance factor Hbs1 (1). GTPBP1/GTPBP2 are encoded by all major groups of eukaryotes except for yeasts and multicellular plants. In many eukaryotes, both GTPBP1 and GTPBP2 representatives are found, which in humans share ∼68% similarity (1–5). Like all other translational GTPases, GTPBP1/GTPBP2 contain three conserved domains—a GTPase (G) domain and two β-barrel domains— but they also have long N-terminal and C-terminal extensions that are the most divergent elements in these proteins (1–4). GTPBP1/GTPBP2 are expressed in a wide range of tissues and organs in mice and in humans, and are localized to the cytoplasm (6, 7).

The loss of either GTPBP1 or GTPBP2 causes neurodegeneration and/or neurodevelopmental disorders in humans (8–12) and in mice that are deficient in the main CNS Arg-tRNA isodecoder (13, 14). The loss of GTPBP1 also disrupts vascular patterning in zebrafish (15). In Drosophila, reduced expression of the GTPBP1/GTPBP2 orthologs Dgp-1 and CG2017 leads to locomotor function defects (12, 16, 17) and neuron-specific repression of Dgp-1 expression results in olfactory learning defects (18).

The molecular function of GTPBP1 is associated with translation and surveillance/quality control mechanisms (14, 19). The studies of neurodegeneration caused by GTPBP1 deletion and Arg-tRNA disruption in mice suggest that GTPBP1 resolves the ribosomes paused during elongation (14). GTPBP1 was also proposed to recruit the exosome to promote mRNA degradation in stalled ribosomal complexes (19, 20). In *Drosophila*, Dgp-1 is upregulated in response to stresses, including oxidative stress (5), thermal stress (21) and hyperoxia (22). Furthermore, Dgp-1 is upregulated in response to over-expression of the endoplasmic reticulum (ER) stress-induced dPERK kinase (23), and loss-of-function alleles of *Drosophila Dgp-1* reduce the viability of Parkin null mutants deficient in Parkin E3 ubiquitin ligase that targets proteins for degradation (24). These observations indicate that GTPBP1 may be protective against stresses affecting translation regulation and proteostasis, in keeping with the protein’s roles in translation and/or quality control.

Biochemical experiments revealed that GTPBP1 possesses eEF1A-like elongation activity and can deliver cognate aminoacyl-tRNA to the ribosomal A site in a GTP-dependent manner (19). However, in contrast to eEF1A, GTP hydrolysis by GTPBP1 was not followed by rapid peptide bond formation, and GTPBP1-mediated elongation was very slow. Additionally, unlike eEF1A, GTPBP1 could also promote ribosomal delivery of deacylated tRNA. Although GTPBP2 was also shown to bind tRNAs in a GTP-dependent manner, it could not promote translation elongation by delivering aminoacyl-tRNA to the ribosome (19).

The structural basis that determines the unusual kinetics of GTPBP1-mediated elongation is unknown. To elucidate the mechanism of GTPBP1 function, we determined cryogenic electron microscopy (cryo-EM) structures of human GTPBP1•aa-tRNA on the 80S ribosome. The intermediates obtained with GTPBP1 and either the non-hydrolysable analog GDPCP or GTP, visualize ribosome complexes prior to or after GTP hydrolysis. They bring insights into the ribosome dynamics, highlighting a role for 40S shoulder domain movements in GTPase activation upon mRNA decoding by GTPBP1•aa-tRNA. These structures reveal how the unique protein architecture of GTPBP1, including an additional N-terminal domain and the shoulder-interacting H-loop in place of the α2 helix of canonical GTPases, is responsible for establishing GTPBP1-specific interactions with tRNA and the ribosome. These interactions are distinct from those of its functional analog eEF1A and account for the prolonged association of GTPBP1•aa-tRNA with the ribosome after GTP hydrolysis and delayed accommodation of tRNA in the A site of the ribosome. Our structures and biochemical data suggest the increased decoding accuracy of GTPBP1, which likely coincides with GTPBP1’s quality-control cellular functions.

## Results

### GTPBP1•aa-tRNA•ribosome complexes in the presence of GDPCP

During canonical mRNA decoding, elongation factors (EF: EF-Tu in bacteria and a/eEF1A in archaea or eukaryotes) bound with GTP and aminoacyl-tRNA sample the ribosomal A site. The EF•GTP•aa-tRNA complex initially binds to the small subunit, placed away from the GTPase-activating sarcin-ricin loop (SRL) of the large subunit (25–27). This placement prevents spontaneous GTP hydrolysis, required for the dissociation of EF from tRNA. If the tRNA anticodon matches the mRNA codon, rearrangements of the small subunit bring the EF closer to the SRL, triggering GTP hydrolysis and EF•GDP dissociation, which allows the accommodation of the tRNA into the A site and formation of a new peptide bond. To visualize the dynamics of GTPBP1 on the ribosome prior to GTP hydrolysis, we took advantage of the non-hydrolysable GTP analog GDPCP (5′-guanosyl-β, γ-methylene-triphosphate), which had allowed visualization of other translational GTPases on the ribosome (28–30). We incubated GTPBP1•Phe-tRNA^GAA^•GDPCP with the 80S ribosomes purified from rabbit reticulocyte lysates and programmed with a derivative of β-globin mRNA and containing initiator Met-tRNA_i_^Met^ bound to the AUG start codon in the P site and the UUC Phe codon in the A-site (19). The sample was subjected to cryo-EM. To elucidate the role of GTP hydrolysis in GTPBP1, we assembled an 80S ribosome complex with GTPBP1•Phe-tRNA^Phe^ and GTP, similarly to the complex described above, placed the mixture on ice and collected two cryo-EM datasets corresponding to 1- and 5-min time points of the reaction.

Classification of cryo-EM data yielded ten maps at average resolutions of 2.9 to 3.1 Å, bringing insights into the stages of mRNA-decoding by GTPBP1: ribosomes with a vacant A site (Structures Ia through Ic featuring slightly different conformations of the 40S subunit, as discussed); ribosomes bound with GTPBP1•tRNA•GDPCP, featuring GTPBP1 away from the GTPase-activating SRL (Structures IIa-d) and GTPBP1 docked at the SRL (Structure III); ribosome bound to GTPBP1•tRNA•GDP (Structure IV); and ribosome with tRNA^Phe^ accommodated in the A-site (Structure V) (Figure 1A, Supplementary Table 1, Supplementary Figure 1).

**Figure 1.**
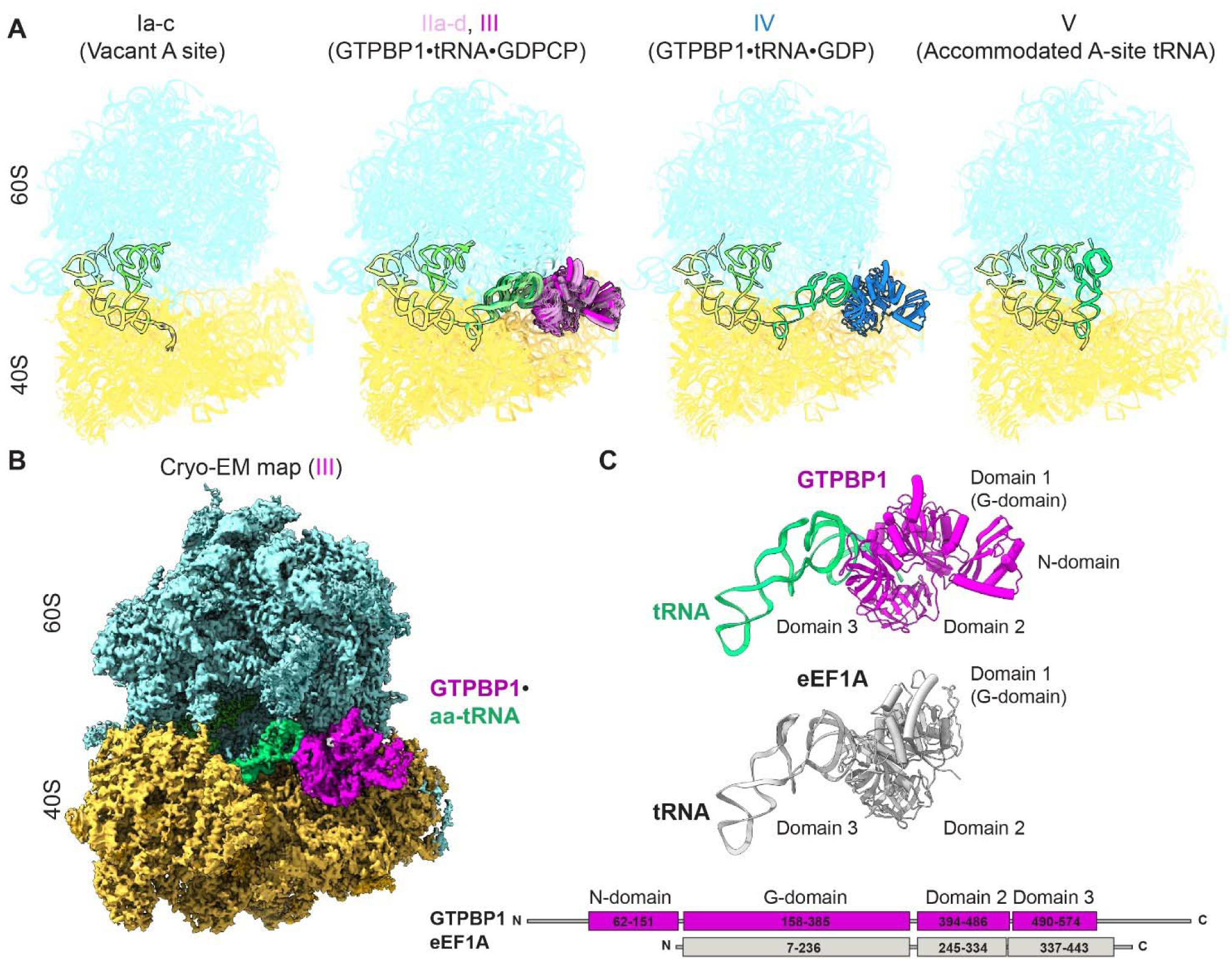
Cryo-EM structures representing. GTPBP1-catalysed decoding on the mammalian 80S ribosome. **A)** The ensemble of models resulted from the analysis of the Cryo-EM data of GTPBP1•GDPCP•aa-tRNA and GTPBP1•GDP•aa-tRNA datasets: Structures Ia-c, featuring a vacant A site; Structures IIa-d (undocked GTPBP1) and III (GTPBP1 docked at SRL), featuring ribosomes with GTPBP1•GDPCP and aa-tRNA; Structure IV (GTPBP1•GDP and aa-tRNA), and Structure V with accommodated A-site aa-tRNA. **B**) Cryo-EM density of ribosome with GTPBP1•GDPCP•aa-tRNA (Structure III, 60S subunit is shown in cyan, 40S subunit yellow, aa-tRNA green, GTPBP1 magenta). **C**) Comparison of domain organizations of human GTPBP1 (in Structure III) and eEF1A (PDB:5LZS).

We first describe the overall structure of GTPBP1, its interactions with tRNA, GTP and the ribosome, and then discuss the structural rearrangements of the ribosome during mRNA decoding. GTPBP1•Phe-tRNA^GAA^•GDPCP binds at the ribosomal A site (Figure 1B). The architecture of GTPBP1 (most well resolved in Structure III, Figure 2B) is similar to that of eEF1A, with the exception of a novel N-terminal domain (aa 62-151), which features two α-helices packed onto a β-sheet, residing on top of the GTPase domain (Figure 1C). Despite the lack of sequence similarity, this domain structurally resembles archaeal/eukaryotic initiation factors e/aIF1 and the C-terminal domain of bacterial initiation factor IF3 (31, 32), according to the DALI fold comparisons (33). The N- and C-termini of GTPBP1 (aa 1-61 and 575-669, respectively), which are predicted to form unfolded tails, are not resolved in our maps, consistent with high conformational flexibility (Figure 1B, C). The two β-barrel domains of GTPBP1 (domains 2 and 3: aa 394-486 and 490-574, respectively) are bound at the shoulder domain of the 40S subunit (Figure 1B, C). Domain 3 binds the acceptor arm of the A/T tRNA (A-site/Ternary complex), while the 3′ CCA end of tRNA is held in the cleft between the G-domain and domain 2, similar to that in eEF1A complexes (Figure 1C) (28, 34).

**Figure 2.**
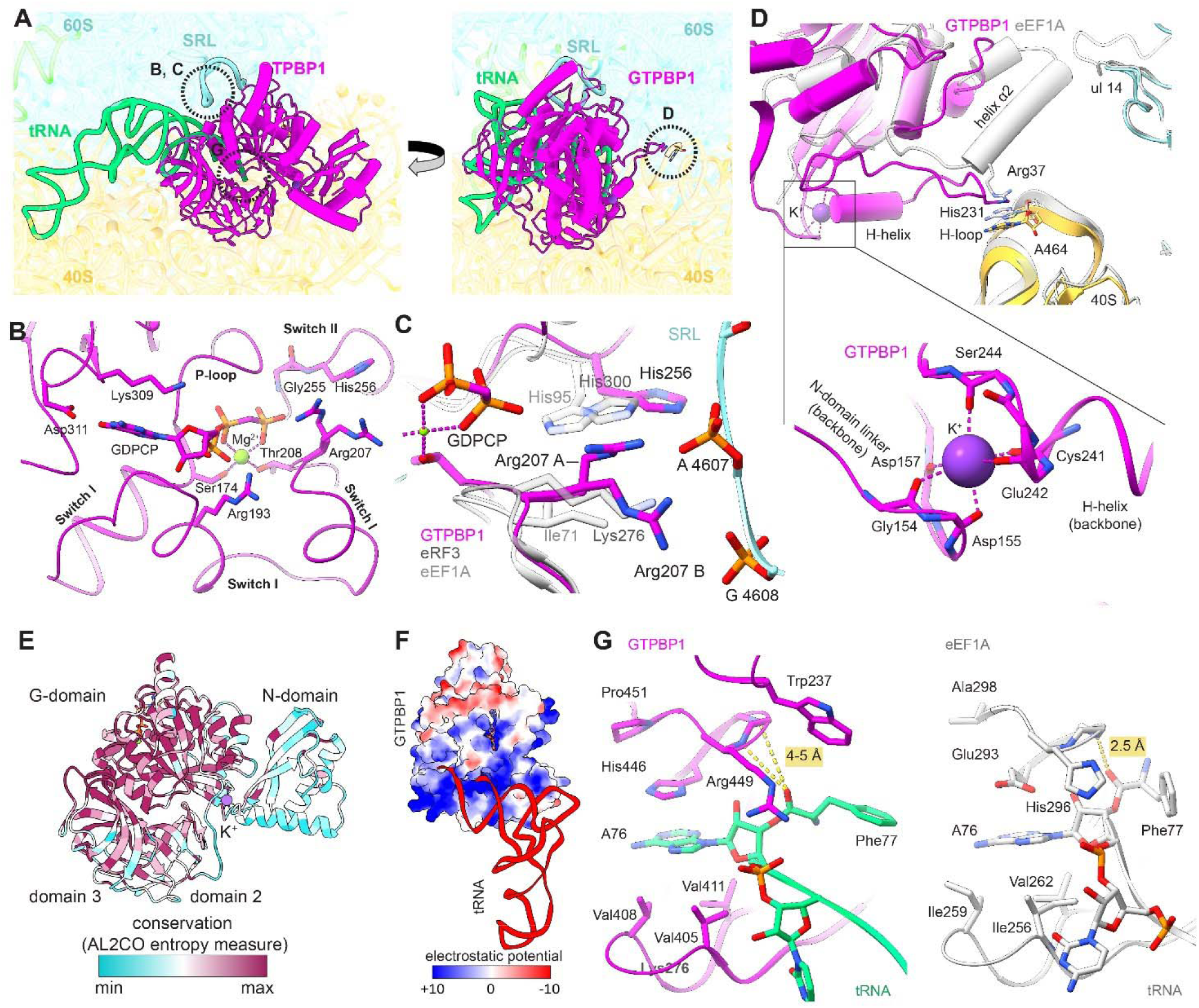
Interactions of GTPBP1 with the ribosome, GDPCP and aa-tRNA (Structure III). **A)** Close-up view of GTPBP1 (magenta) and aa-tRNA (green) on the ribosome. **B)** The nucleotide-binding pocket of GTPBP1. **C)** Comparison of the catalytic GTPase pockets of GTPBP1, eEF1A and eRF3. **D)** Top panel: Distinct interactions of GTPBP1 and eEF1A (5LZS) with 40S and the uL14-mediated bridge between the large and small subunits; the complexes were aligned by 28S rRNA. Lower panel: Potassium-binding site coordinated by the GTPBP1-specific H-helix. **E)** Coloring GTPBP1 according to sequence conservation (39) (sequence alignment is shown in Supplementary Figure 2). **F)** Interactions of GTPBP1 and aa-tRNA, the colored surface represents the electrostatic potential surface of GTPBP1 (red, negative; blue positive). **G)** Interactions of the tip of aa-tRNA with GTPBP1 (left) and eEF1A (8G6J, right).

In the presence of GDPCP mimicking pre-GTP-hydrolysis states, GTPBP1 adopts different positions in the groups of Structures II and Structure III, resolved at ∼3.1 and 2.9 Å average resolutions, respectively (Figures 2-3). Structures IIa through IId differ by slight shifts (∼1 Å) in GTPBP1 and aa-tRNA positions (Figure 1A). In this group of structures, the GTPase domain of GTPBP1 (aa 153-385) is placed ∼7 Å from the SRL (at the catalytic His256 described below) (Figure 3A), resembling a pre-activation state of eEF1A and EF-Tu observed during codon sampling (29, 34–36). The elbow of tRNA (at nucleotides 19 and 56) is stabilized by the 28S rRNA of the L11 stalk at nucleotides 1981 and 2009 (Figure 3C). Unlike the pre-activation state in eEF1A structures (34), where the codon-anticodon interactions are poorly resolved, the anticodon of tRNA is fully paired with the mRNA codon in Structures IIa through IId (Figure 3G).

**Figure 3.**
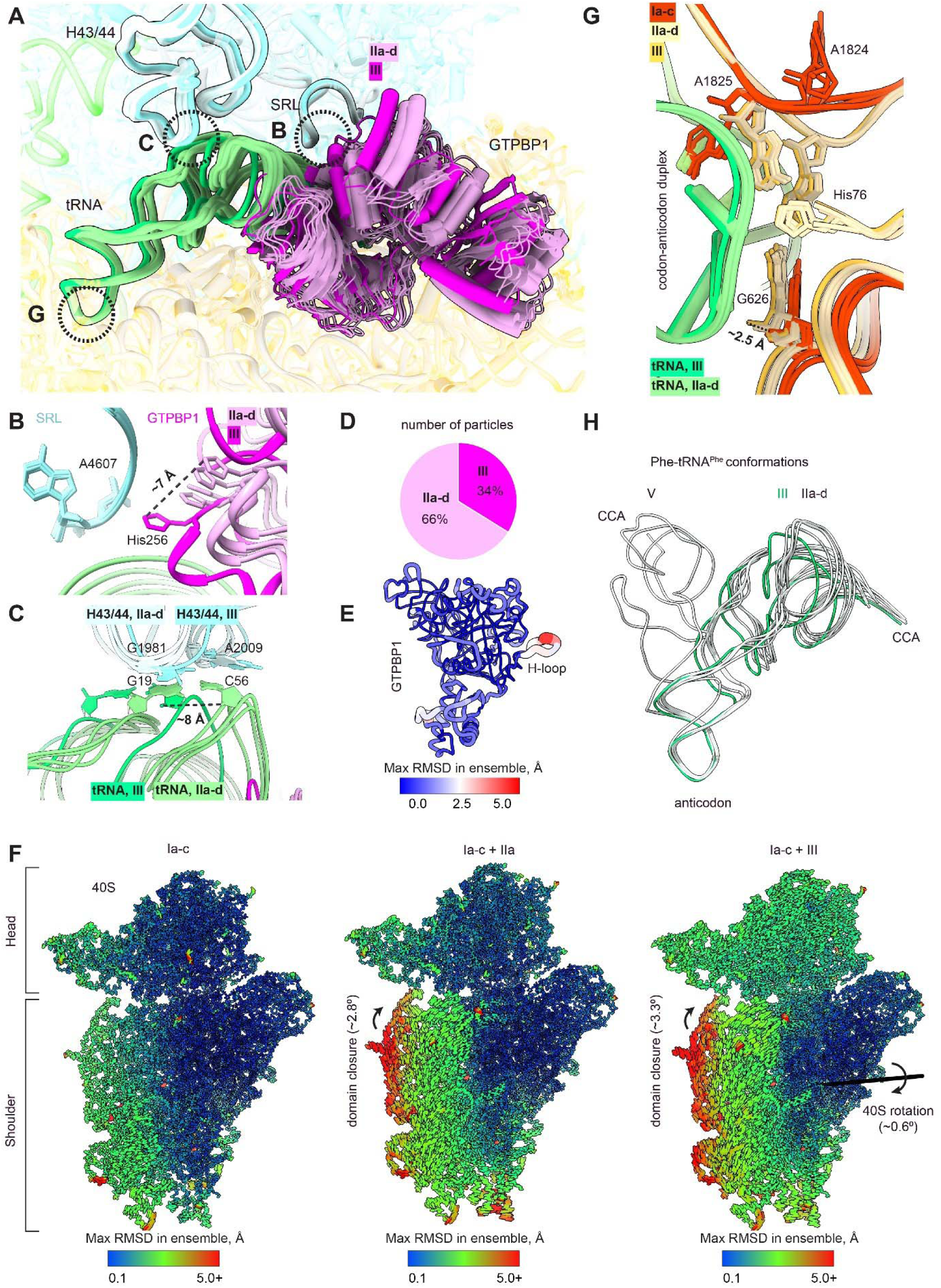
GTPBP1•aa-tRNA•GDPCP adopts different conformations on the ribosome. **A**) Comparison of GTPBP1 positions in Structures IIa-d and III aligned by 28S rRNA. **B**) Positions of catalytic His256 relative to the A4607 of the SRL in Structures IIa-d (pale colors) and III (bright colors). **C**) Comparison of tRNA elbow positions in Structures IIa-d (pale colors) and III (bright colors). **D)** Pie chart representing proportions of ribosome particles corresponding to Structures IIa-d and Structure III. **E**) Worm diagram representing maximal RMSD in the ensemble of GTPBP1•aa-tRNA Structures IIa-d and III aligned via GTPBP1. **F**) Rearrangements of the small subunit of the ribosomes with the vacant A site (Structures Ia-c, left panel) upon binding of GTPBP1•aa-tRNA•GDPCP (Structure IIa, middle panel) and docking of GTPBP1 into the SRL (Structure III, right panel). Cylinders’ lengths and colors are proportional to maximal root-mean-square distances within the ensemble of structures specified on the top of each panel. **G**) Close-up view of the decoding center in Structures IIa-d and III. **H**) Comparison of Phe-tRNA^Phe^ positions in Structures IIa-d through Structure III to Structure V (accommodated tRNA), aligned by 28S rRNA.

In Structure III, GTPBP1 is shifted toward the A site (Figure 2A, 3A), resembling the GTPase activation states of eEF1A and EF-Tu (28, 34). The GTP substrate (mimicked by GDPCP in our structures) is held in place by conserved regions of GTPBP1, including the switch loops (Figure 2B, Supplementary Figure 2a). In translational GTPases, the switch loops must rearrange upon GTP hydrolysis to allow GTPase dissociation from the ribosome (36, 37). The base of GDPCP interacts with the guanine recognition motif [NT]KxD (where x is any amino acid), which in GTPBP1 is represented by ^308^TKID^311^, in contrast to most other translational GTPases featuring NKxD (38). Here, the guanine base is stacked on Lys309 and its Watson-Crick edge interacts with Asp311 (Figure 2B). The triphosphate of GDPCP interacts with the backbone and sidechains of the P-loop (aa 167-174) and Arg193 of switch 1 (aa 182-212). The γ- and β-phosphates are bound to a Mg^2+^ ion coordinated by the conserved Ser174 of the P-loop and Thr208 of switch 1 (Figure 2B). Recently, a T208A mutation was shown to cause neurodegeneration in humans, highlighting the key importance of this residue for GTPBP1 function (12). The γ-phosphate hydrogen bonds with the backbone of the conserved Gly255 in switch 2 (aa 252-269), which is well-resolved in the map next to catalytic His256 (Figure 2B).

The catalytic His256, conserved in translational GTPases (His95 in eEF1A and His300 in eRF3) and required for GTP hydrolysis (28, 34), is placed at the backbone phosphate of A4607 (A2662 in *E. coli*) of the SRL (Figure 2A, C). The side chain of His256 is rotated ∼6 Å from the γ-phosphate (i.e. “outward” conformation), consistent with a catalytically inactive conformation. To allow His256 to move closer, to ∼4 Å from the γ-phosphate (i.e., “inward” conformation), a neighboring residue, Arg207, must shift away. In Structure III, Arg207 appears to adopt two alternative conformations, in which the guanidine group interacts either with the backbone phosphate of A4607 or with both A4607 and G4608 (Figure 2C, Supplementary Figure 3A). The latter rotamer of Arg207 is similar to the position of Lys276 in eRF3, whose catalytic histidine is placed near γ-phosphate (Figure 2C, Supplementary Figure 3A) (28). Thus, the dynamics of GTPBP1 residues His256 and Arg207 appear critical for GTP hydrolysis near the SRL, consistent with the dependence of the GTPase activity on the ribosome (19).

Despite similarity to eEF1A, significant differences in GTPBP1 interactions with its cofactors and the ribosome suggest differences in the dynamics and/or kinetics of GTPBP1-guided tRNA delivery. Perhaps most notable is the absence of helix α2, which in eEF1A and its paralogs eRF3 and Hbs1l is a key part of the switch 1 region, bridging the 40S and 60S subunits during GTPase activation (28, 34). In eEF1A, Arg37 of α2 stacks on A464 at the shoulder of 18S rRNA, while the other end of the helix reaches toward protein uL14 of the 60S subunit (Figure 2D) (28, 34). GTPBP1, however, features a long loop stemming from the GTPase β-sheet that is not part of a switch region (Figure 2D). On the ribosome, the loop is organized into a short β-hairpin (aa 219-236) and a short α-helix (aa 237-243), which resemble a handle and hence are termed the H-hairpin and H-helix (Figure 2D). His231 at the tip of the β-hairpin stacks on A464 of 18S rRNA, likely stabilizing GTPBP1 between the small and large subunits (Figure 2D). Unlike in eEF1A-bound complexes, the N-terminus of uL14 makes no contact with GTPBP1 and is therefore disordered (Figure 2D). Nevertheless, Glu204 and Lys200 of GTPBP1 are positioned to form salt bridges with Lys109 and Glu111 of uL14, resembling the bridging of Glu68 and Lys64 of eEF1A. Thus, in the absence of the rigid α-helix of other GTPases, potentially labile interactions of the ionic groups and dynamic H-loop (see below and Figure 3) of GTPBP1 with the ribosome may confer unique ribosome association and dissociation pathways.

The interface between H-helix and the N-domain linker features strong density within a pocket formed by backbone carbonyl oxygens (Supplementary Figure 3B). Given the characteristic coordination distances of ∼2.8 Å and an octahedral coordination pattern (40–42), the density likely corresponds to a potassium ion (Figure 2D, lower panel). (41). The presence of a K^+^-binding pocket makes GTPBP1 a unique translational GTPase, further suggesting important functional roles for the distinctive H-handle/loop and the N-domain. GTPBP1 dissociation from the tRNA or ribosome may require a rearrangement of the H-handle and/or the N-terminal domain, in which K^+^ exchange may play a role.

Distinct dynamics of GTPBP1 in comparison to eEF1A or EF-Tu are further emphasized by a distinct particle distribution among cryo-EM classes. Unlike eEF1A and EF-Tu, shown by cryo-EM and FRET studies to predominantly adopt a GTPase-activated state with cognate tRNA (29, 34, 36), GTPBP1 is predominantly detached from the SRL, with Structures IIa-d accounting for ∼66% of GTPBP1-bound ribosomes (Figure 3D). Helix α2 of eEF1A was hypothesized to stabilize GTPase-activated state(s) by forming a bridge with both ribosomal subunits (28). The absence of α2 in GTPBP1 therefore may result in GTPBP1 predominantly detached from the large subunit. By contrast, the flexible glycine rich H-loop preserves its contacts with the small subunit in Structures IIa-d and III, helping to accommodate different positions of GTPBP1 (Figure 3E). These differences are likely critical for GTPBP1-mediated accuracy of mRNA decoding as discussed below.

The N-domain, which does not interact with the ribosome or tRNA, contacts the G-domain through hydrophobic and salt bridges interactions (Figure 2E). Beside several residues at the interface, the N-domain has a low sequence conservation (Figure 2E, Supplementary Figure 2a) accompanied by high conservation of the length and spatial organization. This supports an important role of this domain as a structural element, as discussed below.

GTPBP1 interactions with Phe-tRNA^Phe^ reflect the protein’s distinct tRNA substrate preferences and/or tRNA association/dissociation dynamics relative to those of eEF1A. For example, GTPBP1•GTP has high affinities to either amino-acylated tRNAs (Kd of ∼3 nM) and deacylated tRNAs (∼30 nM; (19)). By contrast, eEF1A•GTP strongly prefers aa-tRNA (∼1-5 nM) over deacylated tRNA (up to 10 µM) (43). Whereas the overall pattern of tRNA binding at positively charged interfaces of domain G and β-barrel domains in GTPBP1 is similar to that of eEF1A (Figure 2F), the phenylalanine’s carbonyl attached at the 3′ CCA end of Phe-tRNA^Phe^ (Figure 2G) does not hydrogen-bond with GTPBP1 (4-5 Å), unlike that in the eEF1A complex (2.5 Å; Figure 2G). This likely underlies the less discriminatory binding of GTPBP1 to aa-tRNA and deacyl-tRNAs.

By contrast, the 3′-CCA^76^ end appears to be held more strongly by GTPBP1 than eEF1A. The terminal base of the tRNA is buttressed by a hydrophobic/aromatic stacking system in GTPBP1. Here, the adenine of A76 is sandwiched between the conserved Val405, Val408 and Val411 on one side and His445, in turn stacked on Pro451, on the other side (Figure 2G, Supplementary Figure 3C). This contrasts with eEF1A providing the acidic Glu293 in the place of histidine. The A76 phosphate interacts with Arg449 of GTPBP1, which is packed onto conserved Trp237 of the H-helix of the G-domain (Figure 2G). In eEF1A, His296 is placed similarly to Arg449 to interact with the phosphate group, but the histidine does not appear to contact the G-domain (Figure 2G). Furthermore, the acceptor stem of tRNA at pairs 1:72 and 2:71 is held by a short loop-helix structure (aa 199-205) of the switch 1 region, which must rearrange upon GTP hydrolysis to release a translational GTPase from the ribosome (44). In GTPBP1, the conserved His199 and His201 stack within the tRNA minor groove, while the loop-helix of eEF1A with no aromatic residues appears to have a looser interaction in this region. Overall, the buried surface area between GTPBP1 and aa-tRNA (1655.9 Å^2^) is 15% higher than that between eEF1A and aa-tRNA (1437.6 Å^2^) on the ribosome, suggesting a tighter interaction for GTPBP1. These structural differences may contribute to the distinct dynamics of the tRNA release from GTPBP1 and eEF1A upon GTP hydrolysis, as discussed below.

### Structural insights into GTPase activation

The movement of GTPBP1 relative the SRL (from Structures IIa-d to III) is coupled with changes in tRNA and the 40S subunit (Figure 3), bringing insights into the coordination of codon recognition with GTPase activation. Previous studies showed that in EF-Tu and eEF1 ternary complexes, codon recognition in the ribosomal decoding center of the 40S subunit leads to a closure of the shoulder domain toward the 40S body (28–30, 45). We compared Structures IIa-d and III to Structures Ia-c, which correspond to “pre-delivery” ribosomes, whose A site is vacant and P site is bound with the initiator tRNA (Figure 3F). To distinguish small stochastic movements of the 40S shoulder in three “pre-delivery” maps, we developed “classification of transferred heterogeneity” (CTH; Methods). We find that in “pre-delivery” ribosomes, the shoulder of the 40S subunit undergoes a “rocking” movement within a ∼2-Å range at its periphery (Figures 3F, 3G; Supplementary Video 1). In Structures IIa-d to III, the shoulder progressively shifts by ∼2.5 Å closer to the body than in the ribosomes with a vacant A site (Figures 3F, 3G). Furthermore, the SRL docking of GTPBP1 (Structure IIa-d to III) is accompanied by a subtle SSU rotation (0.5-0.7°), resembling the conformational changes in eEF1A-bound complexes (28, 34) (Figures 3F, 3G; Supplementary Video 1).

Interactions in the decoding center are similar to those with cognate eEF1A complexes (28, 34). As the anticodon loop of Phe-tRNA^GAA^ base-pairs with the UUC codon (Figure 3G), nucleotides A1824 and A1825 (A1492 and A1493 in *E. coli*) flip out of helix 44 of 18S rRNA body to monitor the tRNA-mRNA helix (Figure 3G). They are joined by nucleotide G626 (G530 in bacteria) — at the shoulder domain — which shifts from its position in the “pre-delivery” ribosomes by ∼2.5 Å and, similarly to its bacterial counterpart, rearranges from the *syn* to *anti* conformation (Figure 3G, Supplementary Figure 3D) (29). This range of movement is also observed in bacterial ribosomes with EF-Tu albeit the overall shoulder movement in the 80S ribosome is less pronounced (29, 46). In addition, conserved His76 of eukaryotic protein eS30 becomes resolved to interact with the phosphate of nucleotide A36 of the tRNA anticodon, highlighting a potential contribution of eukaryote-specific proteins in codon recognition (28) (Figure 3G). As the 40S shoulder and GTPBP1 shift toward the SRL, the tRNA elbow slides ∼8 Å along the L11 stalk toward the ribosomal A site, coinciding with the tRNA accommodation trajectory (Figure 3C and 3H).

In summary, our maps demonstrate that similarly to EF-Tu and eEF1A ribosome complexes, the 40S decoding-center rearrangements upon codon-anticodon helix formation are correlated with the 40S shoulder movement toward the body. The movement of the shoulder-bound GTPBP1 brings the GTPase domain toward the SRL for the activation of GTP hydrolysis.

### Ribosome interaction with GTPBP1•aa-tRNA in the presence of GTP

In our previous work, time-resolved cryo-EM allowed us to visualize more than a dozen structural intermediates of mRNA decoding by EF-Tu-delivered tRNA upon GTP hydrolysis (36). GTP hydrolysis leads to the gradual rotation of the G-domain relative to the β-barrel domains (Figure 4F), and similar rearrangements were implicated from the comparison of GTP- and GDP-bound isolated structures of eEF1A (36, 44, 47, 48) and eRF3 (49). The gradual interdomain rearrangement results in the gradual loss of contacts between aa-tRNA and EF-Tu, allowing EF-Tu dissociation and aa-tRNA accommodation into the peptidyl-transferase center (36). Antibiotics kirromycin (for EF-Tu (45)), and didemnin B, plitidepsin and ternatin-4 (for eEF1A (28, 34, 50)) trap these GTPases on the ribosome in the GDP-bound state by binding at the interface between the G domain and second β-barrel, blocking the protein interdomain rearrangements (28, 45).

**Figure 4.**
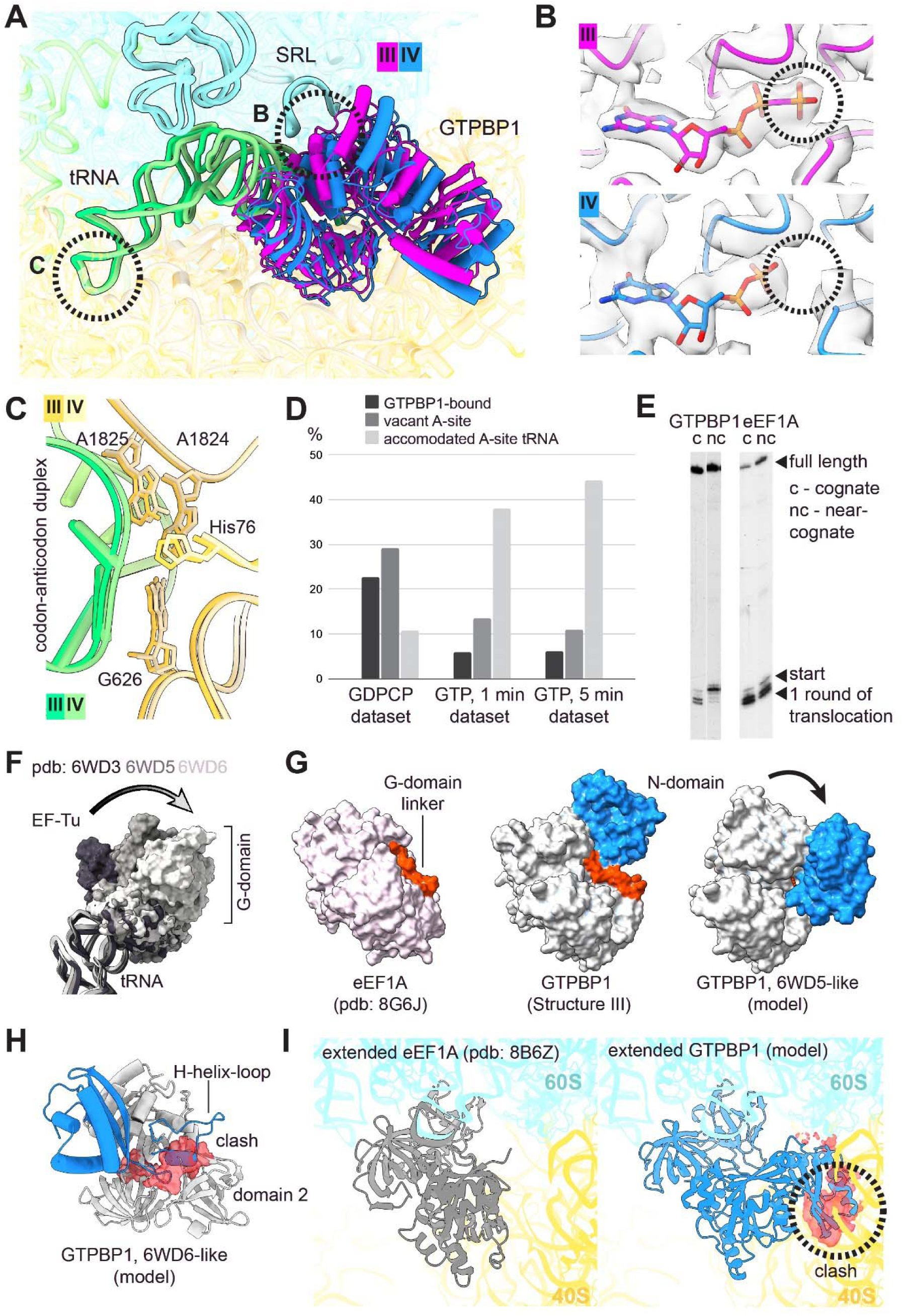
GTPBP1•aa-tRNA•ribosome complexes in the presence of GTP. **A**) Comparison of GTPBP1 position in Structure III and IV aligned by 28S rRNA. **B**) Cryo-EM maps of the nucleotide-binding pockets in complexes Structure III and IV. To improve interpretability of the Structure IV map, the data were refined with a composite mask (51). Black circles indicate location of γ-phosphate. **C**) Close up view of the decoding center in Structure III and IV. **D**) Distribution of particles in datasets obtained in the study. **E**) Toeprinting analysis of the activity of eEF1H and GTPBP1 in one-cycle elongation in the presence of cognate (c) or non-cognate (nc) aa-tRNA. **F**) Structural rearrangements in EF-Tu after GTP-hydrolysis. The models identified on the figure (36) were aligned by domains 2 and 3. **G**) The N-domain can clash with the linker between domains G and 2 after GTP-hydrolysis. The G domain and domains 2+3 of GTPBP1 were separately aligned to the corresponding domains of EF-Tu (6WD5) from (36). **H**) The N-domain linker and H-helix can clash with the domain 2 after GTP-hydrolysis. The G domain and domains 2+3 of GTPBP1 were separately aligned to the corresponding domains of EF-Tu (6WD6) from (36). **I**) The N-domain can interfere with the post-hydrolysis state of GTPBP1 on ribosome. The G-domain of GTPBP1 is aligned against the G-domain of “extended” GDP-bound eEF1A (8B6Z). The rest of ribosome is 5lzs fitted in the map corresponding to 8B6Z (emdb 15893).

Thorough classification of GTP-datasets at 1- and 5-min time points yielded several classes that closely resemble Structures IIa-d with the G-domain of GTPBP1 posed away from the SRL and a codon-anticodon interaction in the decoding center (Figure 4A, C). We term the most well-resolved state Structure IV. We could not identify a sizeable class that resembled the GTPase activation Structure III, with GTPase next to the SRL. This indicates that the GTPase-activated state is only transiently populated by the GTP-bound GTPase, similar to the low abundance “activation” state for EF-Tu•GTP (36).

In contrast to EF-Tu in the previous work, and to GTPBP1 with GDPCP described above, the GTPase center of Structure IV contains GDP (Figure 4B). Gly255 in the switch 2 region, which in GDPCP-bound complexes coordinates the γ-phosphate (Figure 2, 3), is disordered in the presence of GDP. This coincides with the lack of density for the γ-phosphate (Figure 4B), indicating that this region of switch 2 becomes dynamic upon phosphate release, as in other GTPases bound with GDP (50). Despite GTP hydrolysis, GTPBP1 domains retain the conformation observed with GDPCP in Structures IIa-d. The short loop-helix structure of switch 1 (aa 199-205) remains well-resolved in contact with the aa-tRNA, contrasting the rearranged switch 1 initiating EF-Tu•GDP release from the ribosome (36). The GTPBP1’s GTPase domain resembles that in eEF1A•GDP•aa-tRNA complexed with didemnin B, which blocks EF release from the ribosome (28, 34, 50). The inhibition of GTPBP1•GDP rearrangement into an open conformation to release aa-tRNA is likely caused by the extended N-terminal domain, as discussed below. The abundance of ribosome particles with GTPBP1•GDP•aa-tRNA does not change significantly between the two time points (∼5%) (Figure 4D). By contrast, time-resolved cryo-EM showed rapid depletion of ribosome-bound EF-Tu or EF-G over time (36, 37). These results indicate a delayed tRNA accommodation after GTP hydrolysis on GTPBP1, which correlates with the previously observed slow peptide bond formation following rapid hydrolysis of GTP (19). Accordingly, the abundance of ribosomes with accommodated tRNA (collectively in the A/A and post-peptidyl-transfer A/P states) slowly increases from ∼11% in the GDPCP-inhibited complex to 38% and 44% at 1 minutes and 5 minutes, respectively, in the GTP complexes (Figure 4D).

Kinetic proofreading is a key step of aa-tRNA delivery that contributes to the high fidelity of mRNA decoding. While initial selection ensures GTP hydrolysis upon codon-anticodon helix formation, the proofreading step after GTP hydrolysis allows tRNA dissociation to correct for potential initial selection of a near-cognate tRNA (36, 52, 53). A near-cognate tRNA can dissociate together with EF•GDP (54, 55) or alone during accommodation into the peptidyl-transferase center (36, 55, 56). Delayed EF•GDP dissociation must therefore correlate with an increased decoding fidelity (53). Indeed, slow hydrolysis of the GTP analog GTP-γS improves the fidelity of EF-Tu decoding by several orders of magnitude (57). Whether the initial steps of codon recognition are slower for GTPBP1 than for eEF1A remains untested. However, accommodation after GTP-hydrolysis was shown to be slower in biochemical assays (19), which is consistent with our detection of GDP-bound GTPBP1 in Structure IV. These observations suggest that aa-tRNA delivery by GTPBP1 is more accurate than that by eEF1A.

To test this prediction, we assembled mammalian 80S complexes featuring a cognate or a near-cognate codon in the A site, and compared their elongation efficiencies (Methods). Addition of the corresponding aa-tRNA with GTPBP1 or eEF1A, with the translocase eEF2, should result in efficient translocation if the aa-tRNA gets accommodated. Lack of accommodation of a near-cognate aa-tRNA, corresponding to a more accurate decoding, would therefore lead to decreased elongation. Using a toe-printing assay to monitor elongation (19), we found that whereas eEF1A allowed elongation with a near-cognate tRNA, GPTPB1•GTP only allowed elongation with the cognate tRNA (Figure 4E).

What causes the slow dissociation of GTPBP1? Biochemical studies suggested the NTD of GTPBP1 might be important: removing the N-domain (residues 1-152) (Supplementary Figure 4) was shown to increase the rate of aa-tRNA delivery by GTPBP1, resembling that by eEF1A (19). Furthermore, viral GTPBP1-like proteins without the NTD and H-helix-loop drive elongation at rates similar to that of eEF1A (58) (Supplementary Figure 4). Our structural analysis suggests how the unique N-terminal domain residing on top of the GTPase domain may help delay GTPBP1 dissociation from the ribosome. In EF-Tu, the GTPase domain “rotates” away from the tRNA acceptor arm to initiate protein dissociation (Figure 4F) (36). A similar rearrangement in GTPBP1 is likely inhibited by the NTD clash/interaction with the linker connecting the GTPase domain and β-barrel domains (Figure 4G) and/or clash/interaction of the NTD-linker and H-helix-loop with domain 2 (Figure 4H). Similarly, GTPBP1 is not compatible with the recently observed extended conformation of eEF1A bound to the fully accommodated aa-tRNA (35), closely resembling a GDP-bound eEF1A (44). Here, GTPBP1’s NTD would severely clash with the SSU (Figure 4I). These comparisons indicate that the NTD changes the conformational dynamics of GTPBP1, resulting in slower dissociation. The role of the N-domain as a steric block is further supported by its architecture conservation despite high sequence heterogeneity (Figure 2G).

## Discussion

In this study we describe structures of mammalian 80S•GTPBP1•aa-tRNA complexes in the presence of GDPCP and GTP. Although GTPBP1 only shares 14.9% sequence similarity with the canonical elongation factor eEF1A, its structural organization is very similar to that of eEF1A (RMSD of 2.3 Å). The interaction of GTPBP1 with the ribosome also generally resembles that of eEF1A. Nevertheless, our structures uncovered important differences between GTPBP1 and other translational GTPases, yielding mechanistic insights into unique functional aspects of GTPBP1.

The dynamics of GPTBP1 differ from those of eEF1A, appearing to favor conformations with the G-domain posed away from the SRL. Unique structural features of GTPBP1—the N-terminal extension and the shoulder-interacting H-loop in place of canonical α2 helix—underlie interactions of GTPBP1 with the tRNA and the ribosome that are distinct from those of its functional analog eEF1A. Distinct interactions with the ribosome account for the slower dissociation of GTPBP1 compared to other GTPases. Cryo-EM maps reveal that GTP hydrolysis does not quickly lead to the rearrangement of switch 1 of GTPBP1, which is required for the release of aa-tRNA from the protein and aa-tRNA accommodation into the peptidyl-transferase center. Moreover, interdomain rearrangements of GTPBP1 that would be expected to follow GTP hydrolysis are inhibited at least in part by the N-terminal domain. These findings provide mechanistic explanations for the previously reported slow peptide bond formation following GTP hydrolysis by GTPBP1 and accelerated peptide bond formation when the N-terminal domain of GTPBP1 is deleted (19). Indeed, viral GTPBP1-like proteins that lack an NTD and H-helix-loop exhibit high elongation rates similar to those of eEF1A (58). The NTD is connected to the G-domain through a linker coordinating K^+^-binding pocket (Figure 2D). This suggests a possibility of the regulation of the NTD association through cation binding (59), which remains to be tested. In addition, our structures revealed that GTPBP1 interactions with the body of tRNA are tighter than those of eEF1A, perhaps also contributing to tRNA retention and accounting for the unique ability of GTPBP1 to deliver deacylated tRNA to the A site (6).

Because the accuracy of aa-tRNA delivery to the ribosome depends on GTP hydrolysis and aa-tRNA release from a dissociating elongation factor (36, 57), the delayed EF dissociation appears to underlie the more precise aa-tRNA delivery by GTPBP1 than by eEF1A (Figure 4E). GTPBP1 might be required for accurate protein expression at certain stages of cell development or in different cell types, consistent with the differential expression of GTPBP1 in mammalian tissues (2, 7, 14). GTPBP1 is unlikely to compete with the large excess of eEF1A under normal growth conditions, but its elongation function may become more significant under stress or other conditions that impair eEF1A activity. These include eEF1A phosphorylation at Ser300, which inhibits binding of eEF1A to aa-tRNA (60), or phosphorylation of eEF1Bδ at Ser133, which slows nucleotide exchange on eEF1A (61).

Together, our findings suggest the following mechanism of tRNA delivery by GTPBP1•GTP (Figure 5). The ternary complex binds ribosome with GTPBP1 placed on the 40S shoulder away from the GTPase activating SRL. This position allows the anticodon of the tRNA to interact with the mRNA codon in the A site. If the anticodon matches the codon, the decoding center nucleotides G626, A1824 and A1825 stabilize the codon-anticodon helix, inducing a “closure” of the 40S subunit including the movement of the shoulder domain (as in Structure III). The 40S shoulder movement is less extensive than that found in bacterial ribosomes (29, 36) and is similar to that observed in eEF1A-bound complexes (28, 34). In addition, the small ribosomal subunit undergoes a slight <1° forward rotation, which is substantially smaller than the ∼8° rotation observed during tRNA-mRNA translocation (62). Nevertheless, the shoulder movement and the subunit rotation appear sufficient to dock GTPBP1 at the SRL to initiate GTP hydrolysis (as in Structure III). Upon GTP hydrolysis and γ-phosphate release, GTPBP1 undocks from the SRL but preserves its overall conformation and remains bound to the 40S subunit and to the tRNA (Structure IV). The extended residence of the protein on the ribosome likely allows a near-cognate tRNA to dissociate from the ribosome prior to accommodation, accounting for the increased accuracy of decoding (Figure 4E and refs (36, 57)).

**Figure 5.**
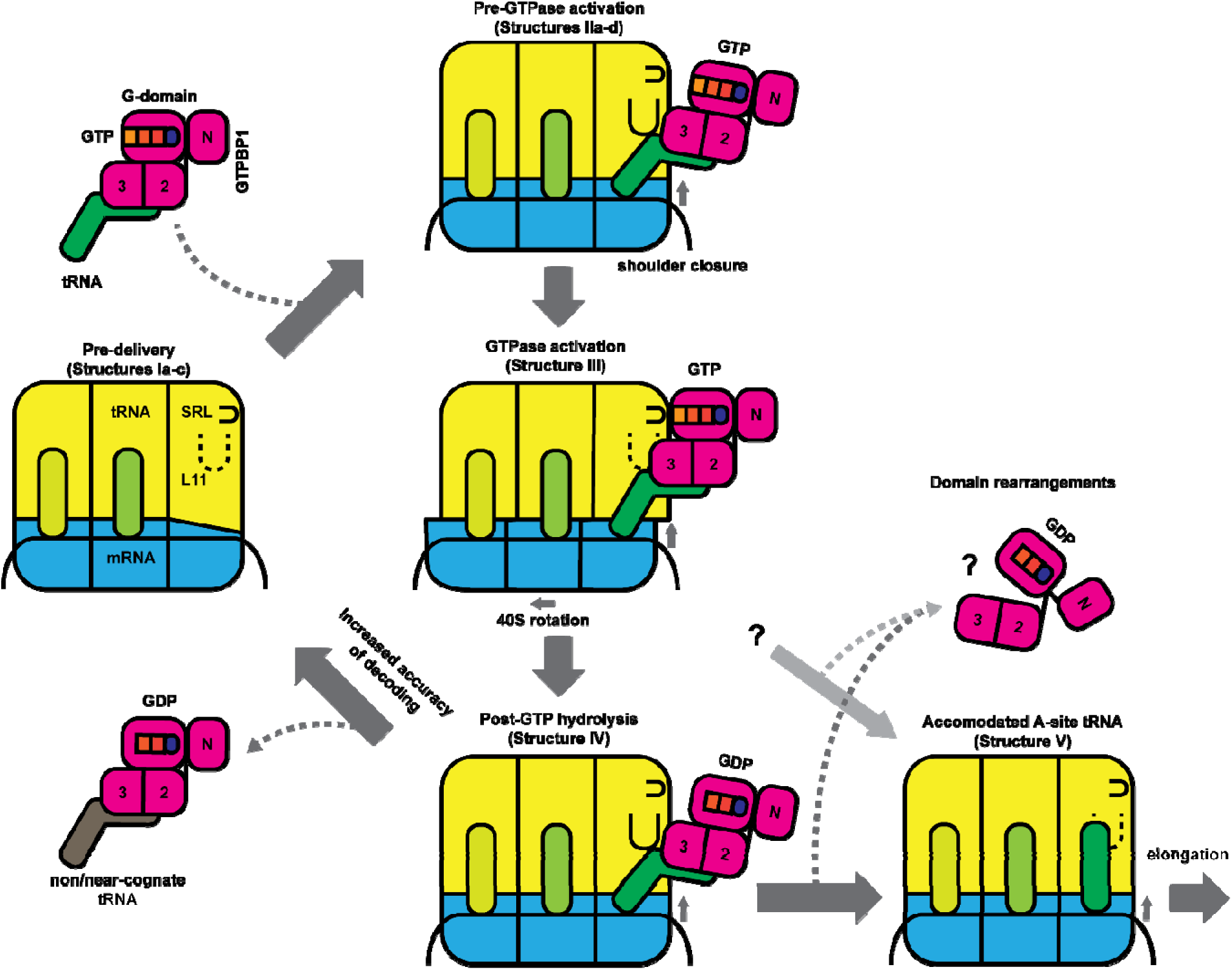
The model of GTPBP1•GTP•aa-tRNA action on the ribosome.

GTPBP1•GDP must eventually rearrange to release aa-tRNA for accommodation and dissociate from the ribosome. Indeed, the fraction of accommodated tRNA is higher when we used GTP for complex assembly than with GDPCP. It remains to be seen how the inhibitory N-terminal domain is housed on the rearranged GTBPB1•GDP. Our alignments suggest that due to steric clashes with the N-terminal domain, the relative positions of the GTPase domain and β-barrel domains likely differ from those in eEF1A.

Our structures also bring insights into GTPBP2, a related protein, which does not bind 80S ribosome under conditions sufficient for GTPBP1 binding (19). Sequence alignment (Supplementary Figure 2b) and AlphaFold prediction of GTPBP2’s structure indicate that GTPBP2 features a substantially shorter H-loop than that of GTPBP1 (Figure 6A, B). The H-loop of GTPBP1 interacts with the 40S subunit and appears to partially replace the conserved α2 helix of eEF1A. Our modeling of GTPBP2 suggests that the protein would be unable to form H-loop-mediated interactions with the 40S subunit. Furthermore, in place of GTPBP1 residues that interact with the small ribosomal subunit, GTPBP2 features numerous negatively charged residues (e.g., at positions 421, 481, 483) that disfavor interactions with the rRNA backbone (Figure 6C, D). Curiously, GTPBP2 retains the N-domain resembling that of GTPBP1, suggesting that the delayed release of its substrate (e.g., tRNA) after GTP-hydrolysis may play a role in GTPBP2 cellular function. Two mutations in this domain of human GTPBP2 (Lys125Arg and Leu93Pro) cause neurodevelopmental impairment and severe intellectual disability (12), in keeping with its functional importance. Deficiencies in GTPBP1 and GTPBP2 similarly affect the CNS of human and tRNA-deficient mice (12–14). However, unlike GTPBP1, GTPBP2 does not seem to interact with the exosome (see below; refs (19, 20)); GTPBP2 and GTPBP1 do not compensate for each other (14); and there is no additive phenotype in the brains of mice lacking both GTPBP1 and GTPBP2 (14). Thus, GTPBP1 and GTPBP2 are thought to function at different steps in the same pathway (8), which would be in line with GTPBP2 performing an enzymatic activity that is distinct from that of GTPBP1.

**Figure 6.**
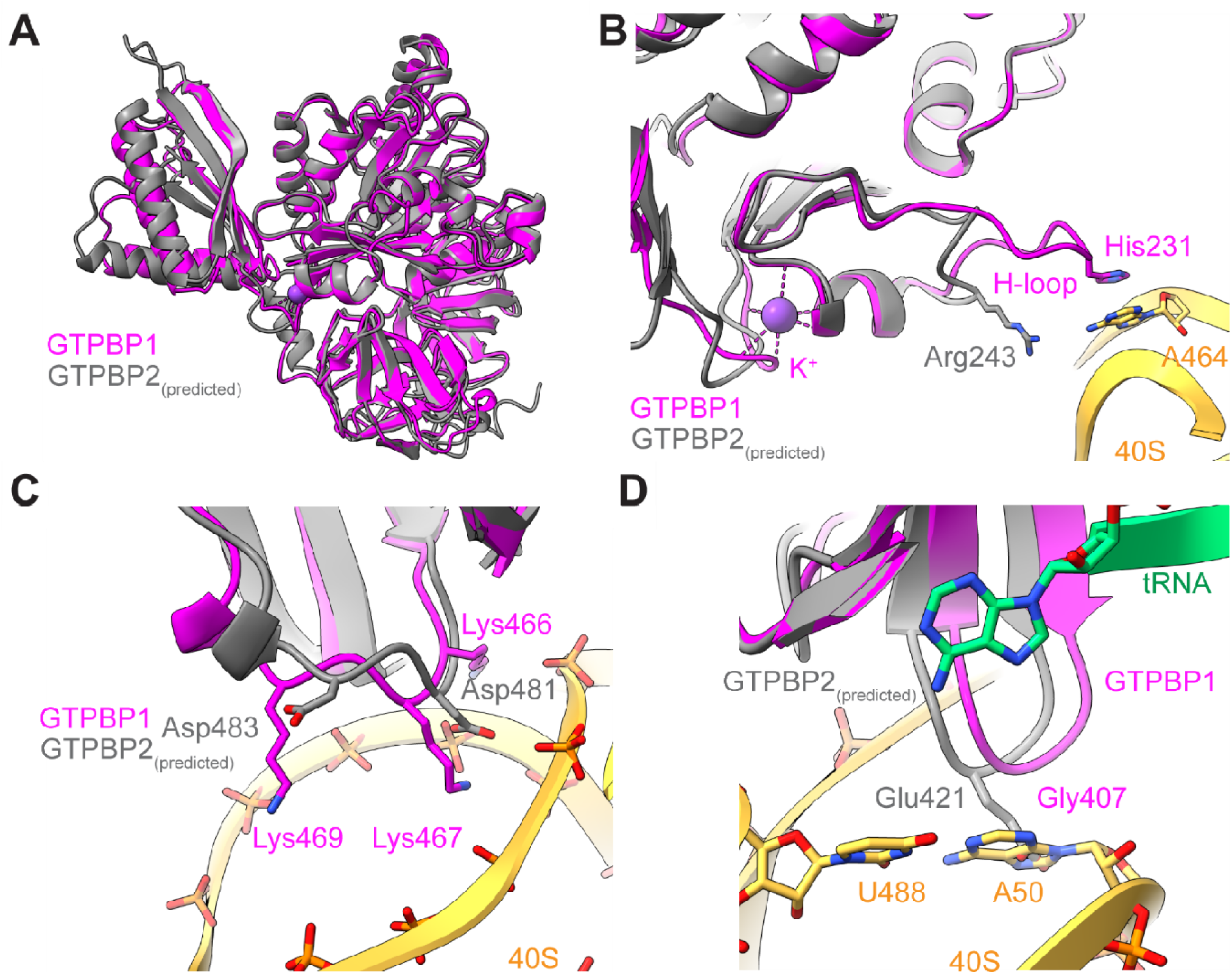
Comparison of GTPBP2 and GTPBP1 on the ribosome. **A**) Overlay of the predicted AlphaFold model of GTPBP2 (Q9BX10) on GTPBP1(Structure III); the long unstructured tails of both proteins are hidden. **B**) H-loop of GTPBP2 is shorter than that of GTPBP1. **C, D**) GTPBP2 bears negatively charged substitutions at points where GTPBP1 contacts ribosome.

It remains to be determined how the molecular mechanism of GTPBP1-mediated translation defines its cellular roles critical for neurodevelopment and stress responses (12, 14, 15). Although GTPBP1 was implicated in the recruitment of the exosome to promote mRNA degradation that could help resolve stalled ribosomes (19, 20, 63), a strong correlation between mRNA abundance and ribosome pausing in the murine *Gtpbp1*^-/-^ cerebellum was not established (8). Alternatively, GTPBP1 could temporarily bind elongating ribosomes with an unclaimed (e.g. “rare”) codon to prevent the formation of ribosomes with a vacant A site that may induce ribosome disassembly (64, 65) and/or the integrated stress response (8). GTPBP1-bound ribosomes might also correspond to a temporary hibernation state, e.g., to enable the dendritic transport of polysomes, in keeping with their stalled or slowed translation (66, 67). Future studies will elucidate the possible interactions of GTPBP1 with other cellular complexes.

In summary, our results provide structural insights into GTPBP1-driven delivery of tRNA to the ribosome, highlighting the similarities with and differences from canonical elongation factors. Similarly to studies of other GTPases (68–70), structures of GTPBP1 may help to develop potential therapies for diseases caused by dysregulation of GTPBP1 function, such as in cancers with altered GTPBP1 expression (71). Structure-based development of GTPBP1 GTPase agonists (72) may help to improve the neurodegenerative conditions caused by GTPBP1 deficiency (12).

## Methods

### Plasmids

Vectors for bacterial expression of His_6_-tagged eIF1, eIF1A (73), eIF4A, eIF4B (74), eIF4G_736-1600_ and eIF4G_736-1115_ (75), eIF5 (76), *Escherichia coli* methionyl-tRNA synthetase (77) and GTPBP1 (19) have been described. Transcription vectors for MF-STOP and MS(UCU)HL-STOP mRNAs containing a 5’UTR derived from the native β-globin mRNA, a short ORF and a 3’UTR composed of nt 16-121 of β-globin ORF have been described (19). The transcription vector for MS(UCC)HL-STOP mRNA had the same composition and was made by GeneWiz, South Plainfield, NJ. Transcription vectors for tRNA^Phe^-GAA (19), tRNA^Ser^-AGA (78), and tRNA_i_^Met^ (79) have been described.

### mRNA and tRNA preparation

Plasmids for transcription of mRNAs were linearized using HindIII (New England Biolabs, Ipswich MA), and plasmids for transcription of tRNA_i_^Met^, tRNA^Phe^-GAA and tRNA^Ser^-AGA were linearized using BstNI (New England Biolabs). All RNAs were transcribed using T7 RNA polymerase (Thermo Scientific) and purified by FPLC using a Superdex 75 Increase 10/300 GL column. tRNA^Phe^-GAA and tRNA^Ser^-AGA were aminoacylated using purified native aminoacyl-tRNA synthetases (80), while tRNA_i_^Met^ was aminoacylated using recombinant *E. coli* methionyl-tRNA synthetase (77).

### Purification of translation factors, ribosomal subunits and aminoacyl-tRNA synthetases

Mammalian native 40S and 60S ribosomal subunits, Σ aminoacyl-tRNA synthetases, eIF2, eIF3, eIF5B, eEF1H and eEF2 were purified from rabbit reticulocyte lysate (RRL) (Green Hectares, Oregon, WI) as described (80, 81). Recombinant His_6_-tagged eIF1, eIF1A, eIF4A, eIF4B, eIF4G_736-1115_, eIF4G_736-1600,_ eIF5 and *E. coli* methionyl tRNA synthetase were expressed in E. coli BL21 (DE3) and purified as described (80).

### Preparation of 80S initiation complexes

80S initiation complexes (ICs) were assembled *in vitro* using individual purified translation components (80). Twelve 200 μl reaction mixtures containing 100 nM 40S subunits, 200 nM MF-STOP mRNA, 270 nM eIF2, 150 nM eIF3, 300 nM eIF1, 300 nM eIF1A, 300 nM eIF4A, 300 nM eIF4B, 150 nM eIF4G_736-1600_ and 200 nM Met-tRNA_i_^Met^ were incubated in buffer A (20 mM Tris, 100 mM KCl, 2.5 mM MgCl_2_, 2 mM DTT, 0.25 mM spermidine, 0.2 mM GTP, 1 mM ATP) for 10 minutes at 37°C. Then reaction mixtures were supplemented with 50 pmol 60S subunits, 100 pmol eIF5, and 100 pmol eIF5B, and incubation continued for another 10 minutes at 37°C. After incubation, reaction mixtures were combined and loaded onto four 10-30% linear sucrose density gradients prepared in buffer B (20 mM Tris pH 7.5, 100 mM KCl, 2.5 mM MgCl_2_, 2 mM DTT and 0.25 mM spermidine) and centrifuged for 1 h 30 minutes at 53, 000 rpm at 4°C in a Beckman SW55Ti rotor. Fractions containing 80S ICs were combined, concentrated to 1.3 pmol/μl on Amicon Ultra ultracel-100K centrifugal filters, transferred to buffer C (20 mM Tris-HCl pH 7.5, 50 mM KCl, 2.5 mM MgCl_2_, 2 mM DTT, 250 mM sucrose) and stored at −80°C.

### Elongation activity of GTPBP1

To compare the abilities of GTPBP1 and eEF1H to promote elongation on cognate and near-cognate codons, elongation with Ser-tRNA^Ser^-AGA was assayed using 80S ICs assembled on MS(UCU)HL-STOP or MS(UCC)HL-STOP mRNA, containing a cognate or near-cognate codon in the A site, respectively. Elongation reactions were assembled essentially as described (19). 48S complexes were formed by incubating 25 nM MS(UCU)HL-STOP or MS(UCC)HL-STOP mRNA with 60 nM 40S subunits, 350 nM eIF1, 350 nM eIF1A, 90 nM eIF2, 60 nM eIF3, 300 nM eIF4A, 60 nM eIF4B, 250 nM eIF4G_736-1115_ and 100 nM Met-tRNA_i_^Met^ in buffer D (20mM Tris-HCl pH 7.5, 3.8 mM MgCl_2_, 100 mM KCl, 0.25 mM spermidine, 2 mM DTT) supplemented with 1 mM ATP, 0.3 mM GTP and 1 U/μl RiboLock RNase inhibitor for 105 minutes at 37°C. To obtain 80S ICs, reaction mixtures were supplemented with 90 nM of 60S subunits, 200 nM eIF5 and 60 nM eIF5B, and incubation continued for an additional 10 min. Elongation was carried out by mixing 80S IC reaction mixtures with 60 nM eEF2, 150 nM eEF1H or GTPBP1, and 300 nM Ser-tRNA^Ser^-AGA, after which incubation continued at 37°C for an additional 15 min. The resulting ribosomal complexes were analyzed by primer extension inhibition (80) using AMV reverse transcriptase (Promega, Madison, WI) and a [^32^P]-labeled primer complementary to nt 149-166 of MS(UCU)HL-STOP and MS(UCC)HL-STOP mRNAs. cDNA products were resolved in 6% polyacrylamide sequencing gels followed by autoradiography.

### Cryo-EM sample preparation

Reaction mixtures containing (final concentrations) 0.56 μM GTPBP1, 1.5 μM *in vitro* transcribed aminoacylated yeast Phe-tRNA^Phe^, 1.6 U/μl Ribolock RNAse inhibitor (Thermo Scientific), 0.2 mM GDPCP or GTP and 1 mM ATP in buffer A were incubated for 5 min at 37°C to allow GTPBP1•GDPCP/GTP•Phe-tRNA^Phe^ complexes to form. Sucrose-density-gradient-purified 80S ICs and Mg(OAc)_2_ were then added to the mixture to 0.13 μM and 5 mM, respectively.

For the GDPCP dataset, the sample was incubated for additional 10 min at 37^0^C and then 4 μl of it were applied to Quantifoil R2/1 holey-carbon grids (EMSDiasum), which were preliminary glow discharged with 20 mA current with negative polarity for 30 s in a PELCO easiGlow glow discharge unit. A Vitrobot Mark IV (ThermoFisher Scientific) was used to plunge-freeze the grids in liquid-nitrogen-cooled liquid ethane. For the GTP datasets, the reaction mixture was placed on ice and 4 μl samples were plunge-frozen at 1- and 5-minute time points, as described above.

### Electron microscopy

Cryo-EM data for 80S•GTPBP1•Phe-tRNA^Phe^ complexes with GDPCP or GTP were collected on a Titan Krios electron microscope (ThermoFisher Scientific) at the UMass Chan Cryo-EM Center, operating at 300 kV and equipped with a Gatan Image Filter (slit width 20 eV) (Gatan Inc.) and a K3 Summit direct electron detector (Gatan Inc.) targeting 0.5–1.5-μm underfocus. SerialEM (82) was used to automatically collect 6703 movies (18 frames, 1.6553 e^-^/Å^2^ per frame, total dose 29.7955 e^-^/Å^2^) for the GDPCP dataset, 14046 movies (20 frames, 1.4877 e^-^/Å^2^ per frame, total dose 29.7531 e^-^/Å^2^) for the GTP-1-min dataset, and 8277 movies (20 frames, 1.5008 e^-^/Å^2^ per frame, total dose 30.0165 e^-^/Å^2^) for the GTP-5-min dataset.

### Data processing

The movies were aligned during data collection (“on the fly”) using IMOD (83) to decompress frames, apply the gain reference, and to correct for image drift and particle damage and bin the super-resolution pixel by 2, yielding 0.83 Å/pixel. The corrected .mrc images were imported into CisTEM (84), and images were selected based on the quality of the CTF fits and image appearances (i.e. the absence of grid damage and crystalline ice deposits) yielding 4455, 13721, and 8097 images for the GDPCP, GTP-1-min, and GTP-5-min datasets, respectively. The raw .tif movies corresponding to the selected micrographs were imported into RELION 4.0 (85, 86) and were motion-corrected using RELION’s implementation of MOTIONCOR2 (87). The aligned images were used for particle picking in crYOLO (88), pretrained on several manually picked images, which yielded 318, 381 particles, 1, 029, 824 particles, and 519, 347 particles for the GDPCP, GTP-1-min, and GTP-5-min datasets, respectively. The particles were extracted in RELION 4.0 at 3.32 Å/pixel in the 140 pixel box and 3D refined using the map of the rabbit unrotated ribosome (emd-9237 (89)), a low-pass filtered to 30 Å, as the initial reference. The resulting map for the GDPCP dataset contained density for GTPBP1, which allowed for initial placing of GTPBP1 model and creating 3D masks around it. The refined stacks were subjected to maximum-likelihood 3D classifications without additional alignments (into 8 classes, Supplementary Figure 5). Classes representing damaged particles, free 60S subunits and rotated ribosomes with hybrid A/P tRNA were discarded. The purified stacks were reextracted at 1.162 Å/pixel, 400-pixel box, and re-refined, followed by cycles of CTF-correction, Bayesian polishing and 3D-refinement. The shiny stacks were exported to Frealign 9.11 (90, 91) and subjected to 3D classifications without alignments with a 3D mask around GTPBP1. The classes containing GTPBP1 were then exported to cryoSPARC v4.3.0 (92) and subjected to 3D classifications without alignments with a 3D mask around GTPBP1. The classes representing different conformations of GTPBP1 relative to the ribosome were imported back into RELION 4.0, re-refined, and the resulting maps were used for structure modelling and refinements. Local resolutions of the maps were estimated with RELION 4.0.

To improve local density at the K^+^-binding pocket and other regions, image stacks were subjected to composite mask refinement (51) in RELION 4.0 using a “protein” mask around GTPBP1 and a “micelle” mask around ribosome and GTPBP1, followed by B-factor sharpening (−50 Å^2^) using bfactor.exe, part of the Frealign v9.11 distribution (90, 91).

In parallel, the initial stacks refined at 3.32 Å/pixel were exported to Frealign 9.11 (90, 91) and subjected to 3D classification into 8 classes (without alignments) with a generous focus mask around the ribosomal A-site for the quantification of the ribosomal states (e.g. A-site occupancies) in the samples.

### The CTH (classification of transferred heterogeneity) approach

We sought to characterize the stochastic movements of the 40S shoulder in 80S ribosomes, but separation of the slightly mobile 40S regions in vacant 80S ribosomes was problematic using standard 3D classifications in RELION 4.0 or Frealign 9.11. To discern small conformational changes, we developed the approach which we term “classification of transferred heterogeneity”, or CTH. It relies on the fact that unlike a 3D classification, particle refinement with a 3D mask covering the shoulder yielded more resolved maps for this region (Supplementary Figure 6). Since the local refinement must “transfer” the local heterogeneity to the rest of ribosome, the heterogeneity at the distant regions must be amplified. Indeed, subsequent classification of the locally refined “pre-delivery” 80S stack, now using a 60S mask, yielded resolved classes with distinct shoulder positions differing by up to 2 Å (Figure 3F, Supplementary Video 1). This approach also identified a small fraction of the A-site accommodated tRNA complexes with the larger shoulder shift due to the domain closure, which was not identified by canonical shoulder-focused classification due to the small number of particles.

### Model building and refinement

The structure of the rabbit 80S complex with eEF1A•GDP•aa-tRNA (28) (PDB: 5LZS) was used as a starting model for the 80S•GTPBP1 complexes. The sequences of mRNA, P-site tRNA and A/T tRNA were changed in Coot (94) to match our mRNA, human Met-tRNA^ini^ and yeast Phe-tRNA^Phe^ sequences, respectively. AlphaFold (95) model of the human GTPBP1 protein (AF-O00178-F1-model_v4 (96)) was used as a starting model for GTPBP1. The structures were assembled in ChimeraX (97), modelled and refined in Isolde (98), followed by B-factor and global minimization refinements in Phenix (99).

### GTPBP1 sequence alignments

For GTPBP1 sequence alignments, we used a set of sequences identified previously to include a broad range of organisms (1). The obsolete records were updated using BlastP (100), and the Ascomycota sequences were excluded as a distant branch of fungi GTPBP1 featuring a different H-loop organization according to AlphaFold (96). Given the absence of the extra N-domain in some GTPBP1-like proteins from giant viruses (58) (Supplementary Figure 4), we checked the remaining sequences using the ChimeraX AlphaFold tool and retained those with the extended N-domain in the predicted models. The resulting 31 unique sequences span from the unicellular amoeboid protist *Capsaspora owczarzaki* to human. GTPBP2 sequences were retrieved from Uniprot (101) using “the species names from the GTPBP1 sequences list AND GTPBP2” as a query, which yielded 15 GTPBP2 sequences. GTPBP1 and GTPBP1/2 sequence sets were aligned using MUSCLE (102) at the EMBL-EBI server (103), and the alignments were visualized in Jalview 2.11.3.3 (104). The model of GTPBP1 was colored by sequence conservation in ChimeraX using GTPBP1 alignment (39).

### Miscellaneous

The structures were visualized and figures panels were prepared using ChimeraX (97). Maximal RMSDs in the ensemble of structures were calculated using a custom Bash script, and ChimeraX was used to visualize the RMSDs as worm or cylinders diagrams. DALI server (33) was used to find structural homologs of the GTPBP1’s N-domain. Figures were assembled in Adobe Illustrator. Video was prepared using ChimeraX and Adobe After Effects. Graphs were prepared in Microsoft Excel.

## Acknowledgements

We thank Chen Xu, Kangkang Song and Christna Ouch for the help with data collection at the cryo-EM facility at UMass Chan Medical School; Darryl Conte Jr., and members of the Pestova and Korostelev laboratories for helpful comments on the manuscript and discussions. This study was supported by grants from the US National Institutes of Health R35GM122602 to T.V.P and R35GM127094 to A.A.K.

## Author contributions

D.S. conceptualized the study, collected and analyzed cryo-EM data, prepared illustrations and video, and wrote the manuscript draft; A.M. and A.Z purified proteins, prepared tRNA and mRNA, assembled 80S complexes and performed biochemical experiments; D.G. and A.B.L. contributed to cryo-EM data analysis; T.V.P. and A.A.K. conceptualized the study, analyzed data and results, wrote the manuscript and secured funding. All authors contributed to data interpretation and provided feedback on the manuscript.

## References

1. Atkinson, C.C. (2015) The evolutionary and functional diversity of classical and lesser-known cytoplasmic and organellar translational GTPases across the tree of life. BMC Genomics, 16, 78.

2. Senju, S. and Nishimura, Y. (1997) Identification of human and mouse GP-1, a putative member of a novel g-protein family. Biochem. Biophys. Res. Commun., 10.1006/bbrc.1997.6103.

3. Kudo, H., Senju, S., Mitsuya, H. and Nishimura, Y. (2000) Mouse and human GTPBP2, newly identified members of the GP-1 family of GTPase. Biochem. Biophys. Res. Commun., 10.1006/bbrc.2000.2763.

4. Watanabe, M., Yoshida, K., Hida, M., Kato, H., Uchida, K., Yamaguchi, R., Tateyama, S. and Sugano, S. (2000) Cloning, expression analysis, and chromosomal mapping of GTPBP2, a novel member of the G protein family. Gene, 10.1016/S0378-1119(00)00346-2.

5. Girardot, F., Monnier, V. and Tricoire, H. (2004) Genome wide analysis of common and specific stress responses in adult drosophila melanogaster. BMC Genomics, 10.1186/1471-2164-5-74.

6. Senju, S., Iyama, K., Kudo, H., Aizawa, S. and Nishimura, Y. (2000) Immunocytochemical Analyses and Targeted Gene Disruption of GTPBP1. Mol. Cell. Biol., 10.1128/mcb.20.17.6195-6200.2000.

7. Anisimova, A.S., Kolyupanova, N.M., Makarova, N.E., Egorov, A.A., Kulakovskiy, I. V. and Dmitriev, S.E. (2023) Human Tissues Exhibit Diverse Composition of Translation Machinery. Int. J. Mol. Sci., 10.3390/ijms24098361.

8. Jaberi, E., Rohani, M., Shahidi, G.A., Nafissi, S., Arefian, E., Soleimani, M., Rasooli, P., Ahmadieh, H., Daftarian, N., Carrami, E.M., et al. (2016) Identification of mutation in GTPBP2 in patients of a family with neurodegeneration accompanied by iron deposition in the brain. Neurobiol. Aging, 10.1016/j.neurobiolaging.2015.10.034.

9. Bertoli-Avella, A.M., Garcia-Aznar, J.M., Brandau, O., Al-Hakami, F., Yüksel, Z., Marais, A., Grüning, N.M., Abbasi Moheb, L., Paknia, O., Alshaikh, N., et al. (2018) Biallelic inactivating variants in the GTPBP2 gene cause a neurodevelopmental disorder with severe intellectual disability. Eur. J. Hum. Genet., 10.1038/s41431-018-0097-3.

10. Carter, M.T., Venkateswaran, S., Shapira-Zaltsberg, G., Davila, J., Humphreys, P., Kernohan, K.D. and Boycott, K.M. (2019) Clinical delineation of GTPBP2-associated neuro-ectodermal syndrome: Report of two new families and review of the literature. Clin. Genet., 10.1111/cge.13523.

11. Manoochehri, J., Shiri, A., Khoddam, S., Aghasipour, M., Kamal, N., Jafari Khamirani, H., Dastgheib, S.A., Dianatpour, M. and Tabei, S.M.B. (2024) Jaberi-Elahi syndrome: Exploring a novel GTPBP2 mutation and a literature review. Eur. J. Med. Genet., 70, 104953.

12. Salpietro, V., Maroofian, R., Zaki, M.S., Wangen, J., Ciolfi, A., Barresi, S., Efthymiou, S., Lamaze, A., Aughey, G.N., Al Mutairi, F., et al. (2024) Bi-allelic genetic variants in the translational GTPases GTPBP1 and GTPBP2 cause a distinct identical neurodevelopmental syndrome. Am. J. Hum. Genet., 111, 200–210.

13. Ishimura, R., Nagy, G., Dotu, I., Zhou, H., Yang, X.L., Schimmel, P., Senju, S., Nishimura, Y., Chuang, J.H. and Ackerman, S.L. (2014) Ribosome stalling induced by mutation of a CNS-specific tRNA causes neurodegeneration. Science (80-.)., 10.1126/science.1249749.

14. Terrey, M., Adamson, S.I., Gibson, A.L., Deng, T., Ishimura, R., Chuang, J.H. and Ackerman, S.L. (2020) GTPBP1 resolves paused ribosomes to maintain neuronal homeostasis. Elife, 10.7554/eLife.62731.

15. Lo, Y.H., Huang, Y.S., Chang, Y.C., Hung, P.Y., Der Wang, W., Liu, W., Urade, R., Wen, Z.H. and Wu, C.Y. (2022) GTP-Binding Protein 1-Like (GTPBP1l) Regulates Vascular Patterning during Zebrafish Development. Biomedicines, 10.3390/biomedicines10123208.

16. Fedotov, S.A., Besedina, N.G., Bragina, J. V., Danilenkova, L. V., Kamysheva, E.A. and Kamyshev, N.G. (2019) Overexpression of isoform B of Dgp-1 gene enhances locomotor activity in senescent Drosophila males and under heat stress. *J. Comp. Physiol. A Neuroethol. Sensory, Neural*, Behav. Physiol., 10.1007/s00359-019-01378-6.

17. Fedotov, S.A., Bragina, J. V., Besedina, N.G., Danilenkova, L. V., Kamysheva, E.A., Panova, A.A. and Kamyshev, N.G. (2014) The effect of neurospecific knockdown of candidate genes for locomotor behavior and sound production in Drosophila melanogaster. Fly (Austin)., 10.4161/19336934.2014.983389.

18. Walkinshaw, E., Gai, Y., Farkas, C., Richter, D., Nicholas, E., Keleman, K. and Davis, R.L. (2015) Identification of genes that promote or inhibit olfactory memory formation in Drosophila. Genetics, 10.1534/genetics.114.173575.

19. Zinoviev, A., Goyal, A., Jindal, S., Lacava, J., Komar, A.A., Rodnina, M. V., Hellen, C.U.T. and Pestova, T. V. (2018) Functions of unconventional mammalian translational GTPases GTPBP1 and GTPBP2. Genes Dev., 10.1101/gad.314724.118.

20. Woo, K., Kim, T., Lee, K., Kim, D., Kim, S., Lee, H., Kang, H., Chung, S.J., Senju, S., Nishimura, Y., et al. (2011) Modulation of exosome-mediated mRNA turnover by interaction of GTP-binding protein 1 (GTPBP1) with its target mRNAs. FASEB J., 10.1096/fj.10-178715.

21. Telonis-Scott, M., van Heerwaarden, B., Johnson, T.K., Hoffmann, A.A. and Sgrò, C.M. (2013) New levels of transcriptome complexity at upper thermal limits in wild Drosophila revealed by exon expression analysis. Genetics, 10.1534/genetics.113.156224.

22. Gruenewald, C., Botella, J.A., Bayersdorfer, F., Navarro, J.A. and Schneuwly, S. (2009) Hyperoxia-induced neurodegeneration as a tool to identify neuroprotective genes in Drosophila melanogaster. Free Radic. Biol. Med., 10.1016/j.freeradbiomed.2009.03.025.

23. Popovic, R., Celardo, I., Yu, Y., Costa, A.C., Loh, S.H.Y. and Martins, L.M. (2021) Combined transcriptomic and proteomic analysis of perk toxicity pathways. Int. J. Mol. Sci., 10.3390/ijms22094598.

24. Greene, J.C., Whitworth, A.J., Andrews, L.A., Parker, T.J. and Pallanck, L.J. (2005) Genetic and genomic studies of Drosophila parkin mutants implicate oxidative stress and innate immune responses in pathogenesis. Hum. Mol. Genet., 10.1093/hmg/ddi074.

25. Korostelev, A.A. (2022) The Structural Dynamics of Translation. Annu. Rev. Biochem., 10.1146/annurev-biochem-071921-122857.

26. Rodnina, M. V. (2023) Decoding and Recoding of mRNA Sequences by the Ribosome. Annu. Rev. Biophys., 10.1146/annurev-biophys-101922-072452.

27. Prabhakar, A., Puglisi, E.V. and Puglisi, J.D. (2019) Single-molecule fluorescence applied to translation. Cold Spring Harb. Perspect. Biol., 10.1101/cshperspect.a032714.

28. Shao, S., Murray, J., Brown, A., Taunton, J., Ramakrishnan, V. and Hegde, R.S. (2016) Decoding Mammalian Ribosome-mRNA States by Translational GTPase Complexes. Cell, 167, 1229–1240.e15.

29. Loveland, A.B., Demo, G., Grigorieff, N. and Korostelev, A.A. (2017) Ensemble cryo-EM elucidates the mechanism of translation fidelity. Nature, 546, 113–117.

30. Voorhees, R.M., Schmeing, T.M., Kelley, A.C. and Ramakrishnan, V. (2010) The mechanism for activation of GTP hydrolysis on the ribosome. Science (80-.)., 330, 835–838.

31. Biou, V., Shu, F. and Ramakrishnan, V. (1995) X-ray crystallography shows that translational initiation factor IF3 consists of two compact α/β domains linked by an α-helix. EMBO J., 10.1002/j.1460-2075.1995.tb00077.x.

32. Fletcher, C.M., Pestova, T. V., Hellen, C.U.T. and Wagner, G. (1999) Structure and interactions of the translation initiation factor eIF1. EMBO J., 10.1093/emboj/18.9.2631.

33. Holm, L. (2020) DALI and the persistence of protein shape. Protein Sci., 10.1002/pro.3749.

34. Holm, M., Natchiar, S.K., Rundlet, E.J., Myasnikov, A.G., Watson, Z.L., Altman, R.B., Wang, H.Y., Taunton, J. and Blanchard, S.C. (2023) mRNA decoding in human is kinetically and structurally distinct from bacteria. Nature, 10.1038/s41586-023-05908-w.

35. Gemmer, M., Chaillet, M.L., van Loenhout, J., Cuevas Arenas, R., Vismpas, D., Gröllers-Mulderij, M., Koh, F.A., Albanese, P., Scheltema, R.A., Howes, S.C., et al. (2023) Visualization of translation and protein biogenesis at the ER membrane. Nature, 10.1038/s41586-022-05638-5.

36. Loveland, A.B., Demo, G. and Korostelev, A.A. (2020) Cryo-EM of elongating ribosome with EF-Tu•GTP elucidates tRNA proofreading. Nature, 584, 640–645.

37. Carbone, C.E., Loveland, A.B., Gamper, H.B., Hou, Y.M., Demo, G. and Korostelev, A.A. (2021) Time-resolved cryo-EM visualizes ribosomal translocation with EF-G and GTP. Nat. Commun., 10.1038/s41467-021-27415-0.

38. Leipe, D.D., Wolf, Y.I., Koonin, E. V. and Aravind, L. (2002) Classification and evolution of P-loop GTPases and related ATPases. J. Mol. Biol., 10.1006/jmbi.2001.5378.

39. Pei, J. and Grishin, N. V. (2001) AL2CO: Calculation of positional conservation in a protein sequence alignment. Bioinformatics, 10.1093/bioinformatics/17.8.700.

40. Zheng, H., Cooper, D.R., Porebski, P.J., Shabalin, I.G., Handing, K.B. and Minor, W. (2017) CheckMyMetal: A macromolecular metal-binding validation tool. Acta Crystallogr. Sect. D Struct. Biol., 10.1107/S2059798317001061.

41. Rozov, A., Khusainov, I., El Omari, K., Duman, R., Mykhaylyk, V., Yusupov, M., Westhof, E., Wagner, A. and Yusupova, G. (2019) Importance of potassium ions for ribosome structure and function revealed by long-wavelength X-ray diffraction. Nat. Commun., 10.1038/s41467-019-10409-4.

42. Auffinger, P., Ennifar, E. and D’Ascenzo, L. (2021) Deflating the RNA Mg2+bubble: Stereochemistry to the rescue! RNA, 10.1261/rna.076067.120.

43. Dreher, T.W., Uhlenbeck, O.C. and Browning, K.S. (1999) Quantitative assessment of EF-1α · GTP binding to aminoacyl-tRNAs, aminoacyl-viral RNA, and tRNA shows close correspondence to the RNA binding properties of EF-Tu. J. Biol. Chem., 10.1074/jbc.274.2.666.

44. Crepin, T., Shalak, V.F., Yaremchuk, A.D., Vlasenko, D.O., McCarthy, A., Negrutskii, B.S., Tukalo, M.A. and El’skaya, A. V. (2014) Mammalian translation elongation factor eEF1A2: X-ray structure and new features of GDP/GTP exchange mechanism in higher eukaryotes. Nucleic Acids Res., 10.1093/nar/gku974.

45. Martin Schmeing, T., Voorhees, R.M., Kelley, A.C., Gao, Y.G., Murphy IV, F. V., Weir, J.R. and Ramakrishnan, V. (2009) The crystal structure of the ribosome bound to EF-Tu and aminoacyl-tRNA. Science (80-.)., 10.1126/science.1179700.

46. Fislage, M., Zhang, J., Brown, Z.P., Mandava, C.S., Sanyal, S., Ehrenberg, M. and Frank, J. (2018) Cryo-EM shows stages of initial codon selection on the ribosome by aa-tRNA in ternary complex with GTP and the GTPase-deficient EF-TuH84A. Nucleic Acids Res., 10.1093/nar/gky346.

47. Maruyama, K., Imai, H., Kawamura, M., Ishino, S., Ishino, Y., Ito, K. and Uchiumi, T. (2019) Switch of the interactions between the ribosomal stalk and EF1A in the GTP- and GDP-bound conformations. Sci. Rep., 10.1038/s41598-019-51266-x.

48. Ito, K., Honda, T., Suzuki, T., Miyoshi, T., Murakami, R., Yao, M. and Uchiumi, T. (2014) Molecular insights into the interaction of the ribosomal stalk protein with elongation factor 1α. Nucleic Acids Res., 10.1093/nar/gku1248.

49. Kong, C., Ito, K., Walsh, M.A., Wada, M., Liu, Y., Kumar, S., Barford, D., Nakamura, Y. and Song, H. (2004) Crystal structure and functional analysis of the eukaryotic class II release factor eRF3 from S. pombe. Mol. Cell, 10.1016/S1097-2765(04)00206-0.

50. Juette, M.F., Carelli, J.D., Rundlet, E.J., Brown, A., Shao, S., Ferguson, A., Wasserman, M.R., Holm, M., Taunton, J. and Blanchard, S.C. (2022) Didemnin B and ternatin-4 differentially inhibit conformational changes in eEF1A required for aminoacyl-tRNA accommodation into mammalian ribosomes. Elife, 10.7554/eLife.81608.

51. Dou, T., Lian, T., Shu, S., He, Y. and Jiang, J. (2023) The substrate and inhibitor binding mechanism of polyspecific transporter OAT1 revealed by high-resolution cryo-EM. Nat. Struct. Mol. Biol., 10.1038/s41594-023-01123-3.

52. Hopefield, J.J. (1974) Kinetic proofreading: a new mechanism for reducing errors in biosynthetic processes requiring high specificity. Proc. Natl. Acad. Sci. U. S. A., 10.1073/pnas.71.10.4135.

53. Zaher, H.S. and Green, R. (2009) Fidelity at the Molecular Level: Lessons from Protein Synthesis. Cell, 10.1016/j.cell.2009.01.036.

54. Zhang, J., Ieong, K.W., Johansson, M. and Ehrenberg, M. (2015) Accuracy of initial codon selection by aminoacyl-tRNAs on the mRNA-programmed bacterial ribosome. Proc. Natl. Acad. Sci. U. S. A., 10.1073/pnas.1506823112.

55. Gromadski, K.B. and Rodnina, M. V. (2004) Kinetic Determinants of High-Fidelity tRNA Discrimination on the Ribosome. Mol. Cell, 10.1016/S1097-2765(04)00005-X.

56. Gavrilova, L.P., Perminova, I.N. and Spirin, A.S. (1981) Elongation factor Tu can reduce translation errors in poly(U)-directed cell-free systems. J. Mol. Biol., 10.1016/0022-2836(81)90260-6.

57. Thompson, R.C. and Karim, A.M. (1982) The accuracy of protein biosynthesis is limited by its speed: High fidelity selection by ribosomes of aminoacyl-tRNA ternary complexes containing GTP[γS]. Proc. Natl. Acad. Sci. U. S. A., 10.1073/pnas.79.16.4922.

58. Zinoviev, A., Kuroha, K., Pestova, T. V. and Hellen, C.U.T. (2019) Two classes of EF1-family translational GTPases encoded by giant viruses. Nucleic Acids Res., 10.1093/nar/gkz296.

59. Page, M.J. and Di Cera, E. (2006) Role of Na+ and K+ in enzyme function. Physiol. Rev., 10.1152/physrev.00008.2006.

60. Lin, K.W., Yakymovych, I., Jia, M., Yakymovych, M. and Souchelnytskyi, S. (2010) Phosphorylation of eEF1A1 at Ser300 by TβR-I results in inhibition of mRNA translation. Curr. Biol., 10.1016/j.cub.2010.08.017.

61. Sivan, G., Aviner, R. and Elroy-Stein, O. (2011) Mitotic modulation of translation elongation factor 1 leads to hindered tRNA delivery to ribosomes. J. Biol. Chem., 10.1074/jbc.M111.255810.

62. Flis, J., Holm, M., Rundlet, E.J., Loerke, J., Hilal, T., Dabrowski, M., Bürger, J., Mielke, T., Blanchard, S.C., Spahn, C.M.T., et al. (2018) tRNA Translocation by the Eukaryotic 80S Ribosome and the Impact of GTP Hydrolysis. Cell Rep., 10.1016/j.celrep.2018.11.040.

63. Zinoviev, A., Hellen, C.U.T. and Pestova, T. V. (2020) In Vitro Characterization of the Activity of the Mammalian RNA Exosome on mRNAs in Ribosomal Translation Complexes. Methods Mol. Biol., 10.1007/978-1-4939-9822-7_16.

64. Filbeck, S., Cerullo, F., Pfeffer, S. and Joazeiro, C.A.P. (2022) Ribosome-associated quality-control mechanisms from bacteria to humans. Mol. Cell, 10.1016/j.molcel.2022.03.038.

65. Inada, T. and Beckmann, R. (2024) Mechanisms of Translation-coupled Quality Control. J. Mol. Biol., 10.1016/j.jmb.2024.168496.

66. Langille, J.J., Ginzberg, K. and Sossin, W.S. (2019) Polysomes identified by live imaging of nascent peptides are stalled in hippocampal and cortical neurites. Learn. Mem., 10.1101/lm.049965.119.

67. Dalla Costa, I., Buchanan, C.N., Zdradzinski, M.D., Sahoo, P.K., Smith, T.P., Thames, E., Kar, A.N. and Twiss, J.L. (2021) The functional organization of axonal mRNA transport and translation. Nat. Rev. Neurosci., 10.1038/s41583-020-00407-7.

68. Lu, S., Jang, H., Gu, S., Zhang, J. and Nussinov, R. (2016) Drugging Ras GTPase: A comprehensive mechanistic and signaling structural view. Chem. Soc. Rev., 10.1039/c5cs00911a.

69. Zhang, H., Cai, J., Yu, S., Sun, B. and Zhang, W. (2023) Anticancer Small-Molecule Agents Targeting Eukaryotic Elongation Factor 1A: State of the Art. Int. J. Mol. Sci., 10.3390/ijms24065184.

70. Gao, Y., Dickerson, J.B., Guo, F., Zheng, J. and Zheng, Y. (2004) Rational design and characterization of a Rac GTPase-specific small molecule inhibitor. Proc. Natl. Acad. Sci. U. S. A., 10.1073/pnas.0307512101.

71. Hu, Y., Chen, L., Tang, Q., Wei, W., Cao, Y., Xie, J. and Ji, J. (2022) Pan-cancer analysis revealed the significance of the GTPBP family in cancer. Aging (Albany. NY)., 10.18632/aging.203952.

72. Palsuledesai, C.C., Surviladze, Z., Waller, A., Miscioscia, T.F., Guo, Y., Wu, Y., Strouse, J., Romero, E., Salas, V.M., Curpan, R., et al. (2018) Activation of Rho Family GTPases by Small Molecules. ACS Chem. Biol., 10.1021/acschembio.8b00038.

73. Pestova, T. V., Borukhov, S.I. and Hellen, C.U.T. (1998) Eukaryotic ribosomes require initiation factors 1 and 1A to locate initiation codons. Nature, 10.1038/29703.

74. Pestova, T. V., Hellen, C.U.T. and Shatsky, I.N. (1996) Canonical Eukaryotic Initiation Factors Determine Initiation of Translation by Internal Ribosomal Entry. Mol. Cell. Biol., 10.1128/mcb.16.12.6859.

75. Lomakin, I.B., Hellen, C.U.T. and Pestova, T. V. (2000) Physical Association of Eukaryotic Initiation Factor 4G (eIF4G) with eIF4A Strongly Enhances Binding of eIF4G to the Internal Ribosomal Entry Site of Encephalomyocarditis Virus and Is Required for Internal Initiation of Translation. Mol. Cell. Biol., 10.1128/mcb.20.16.6019-6029.2000.

76. Pestova, T. V., Lomakin, I.B., Lee, J.H., Choi, S.K., Dever, T.E. and Hellen, C.U.T. (2000) The joining of ribosomal subunits in eukaryotes requires eIF5B. Nature, 10.1038/35002118.

77. Lomakin, I.B., Shirokikh, N.E., Yusupov, M.M., Hellen, C.U.T. and Pestova, T. V. (2006) The fidelity of translation initiation: Reciprocal activities of elF1, IF3 and YciH. EMBO J., 10.1038/sj.emboj.7600904.

78. Zinoviev, A., Hellen, C.U.T. and Pestova, T. V. (2015) Multiple Mechanisms of Reinitiation on Bicistronic Calicivirus mRNAs. Mol. Cell, 10.1016/j.molcel.2015.01.039.

79. Pestova, T. and Hellen, C.U.T. (2001) Preparation and activity of synthetic unmodified mammalian tRNAiMet in initiation of translation in vitro. RNA, 10.1017/S135583820101038X.

80. Pisarev, A. V., Unbehaun, A., Hellen, C.U.T. and Pestova, T. V. (2007) Assembly and Analysis of Eukaryotic Translation Initiation Complexes. In Methods in Enzymology.

81. Pestova, T. V. and Hellen, C.U.T. (2003) Translation elongation after assembly of ribosomes on the Cricket paralysis virus internal ribosomal entry site without initiation factors or initiator tRNA. Genes Dev., 17, 181–186.

82. Mastronarde, D.N. (2005) Automated electron microscope tomography using robust prediction of specimen movements. J. Struct. Biol., 10.1016/j.jsb.2005.07.007.

83. Kremer, J.R., Mastronarde, D.N. and McIntosh, J.R. (1996) Computer visualization of three-dimensional image data using IMOD. J. Struct. Biol., 10.1006/jsbi.1996.0013.

84. Grant, T., Rohou, A. and Grigorieff, N. (2018) CisTEM, user-friendly software for single-particle image processing. Elife, 7.

85. Kimanius, D., Dong, L., Sharov, G., Nakane, T. and Scheres, S.H.W. (2021) New tools for automated cryo-EM single-particle analysis in RELION-4.0. Biochem. J., 10.1042/BCJ20210708.

86. Scheres, S.H.W. (2012) RELION: Implementation of a Bayesian approach to cryo-EM structure determination. J. Struct. Biol., 10.1016/j.jsb.2012.09.006.

87. Zheng, S.Q., Palovcak, E., Armache, J.P., Verba, K.A., Cheng, Y. and Agard, D.A. (2017) MotionCor2: Anisotropic correction of beam-induced motion for improved cryo-electron microscopy. Nat. Methods, 10.1038/nmeth.4193.

88. Wagner, T., Merino, F., Stabrin, M., Moriya, T., Antoni, C., Apelbaum, A., Hagel, P., Sitsel, O., Raisch, T., Prumbaum, D., et al. (2019) SPHIRE-crYOLO is a fast and accurate fully automated particle picker for cryo-EM. *Commun*. Biol., 10.1038/s42003-019-0437-z.

89. Brown, A., Baird, M.R., Yip, M.C.J., Murray, J. and Shao, S. (2018) Structures of translationally inactive mammalian ribosomes. Elife, 10.7554/eLife.40486.

90. Grigorieff, N. (2016) Frealign: An Exploratory Tool for Single-Particle Cryo-EM. In Methods in Enzymology.Vol. 579, pp. 191–226.

91. Lyumkis, D., Brilot, A.F., Theobald, D.L. and Grigorieff, N. (2013) Likelihood-based classification of cryo-EM images using FREALIGN. J. Struct. Biol., 10.1016/j.jsb.2013.07.005.

92. Punjani, A., Rubinstein, J.L., Fleet, D.J. and Brubaker, M.A. (2017) CryoSPARC: Algorithms for rapid unsupervised cryo-EM structure determination. Nat. Methods, 10.1038/nmeth.4169.

93. Heymann, J.B. (2022) Bsoft: Image Processing for Structural Biology. Bio-protocol, 10.21769/BioProtoc.4393.

94. Emsley, P., Lohkamp, B., Scott, W.G. and Cowtan, K. (2010) Features and development of Coot. Acta Crystallogr. Sect. D Biol. Crystallogr., 10.1107/S0907444910007493.

95. Jumper, J., Evans, R., Pritzel, A., Green, T., Figurnov, M., Ronneberger, O., Tunyasuvunakool, K., Bates, R., Žídek, A., Potapenko, A., et al. (2021) Highly accurate protein structure prediction with AlphaFold. Nature, 10.1038/s41586-021-03819-2.

96. Varadi, M., Bertoni, D., Magana, P., Paramval, U., Pidruchna, I., Radhakrishnan, M., Tsenkov, M., Nair, S., Mirdita, M., Yeo, J., et al. (2024) AlphaFold Protein Structure Database in 2024: providing structure coverage for over 214 million protein sequences. Nucleic Acids Res., 10.1093/nar/gkad1011.

97. Meng, E.C., Goddard, T.D., Pettersen, E.F., Couch, G.S., Pearson, Z.J., Morris, J.H. and Ferrin, T.E. (2023) UCSF ChimeraX: Tools for structure building and analysis. Protein Sci., 10.1002/pro.4792.

98. Croll, T.I. (2018) ISOLDE: A physically realistic environment for model building into low-resolution electron-density maps. Acta Crystallogr. Sect. D Struct. Biol., 10.1107/S2059798318002425.

99. Liebschner, D., Afonine, P. V., Baker, M.L., Bunkoczi, G., Chen, V.B., Croll, T.I., Hintze, B., Hung, L.W., Jain, S., McCoy, A.J., et al. (2019) Macromolecular structure determination using X-rays, neutrons and electrons: Recent developments in Phenix. Acta Crystallogr. Sect. D Struct. Biol., 10.1107/S2059798319011471.

100. Altschul, S.F., Gish, W., Miller, W., Myers, E.W. and Lipman, D.J. (1990) Basic local alignment search tool. J. Mol. Biol., 10.1016/S0022-2836(05)80360-2.

101. Bateman, A., Martin, M.J., Orchard, S., Magrane, M., Ahmad, S., Alpi, E., Bowler-Barnett, E.H., Britto, R., Bye-A-Jee, H., Cukura, A., et al. (2023) UniProt: the Universal Protein Knowledgebase in 2023. Nucleic Acids Res., 10.1093/nar/gkac1052.

102. Edgar, R.C. (2004) MUSCLE: Multiple sequence alignment with high accuracy and high throughput. Nucleic Acids Res., 10.1093/nar/gkh340.

103. Madeira, F., Madhusoodanan, N., Lee, J., Eusebi, A., Niewielska, A., Tivey, A.R.N., Lopez, R. and Butcher, S. (2024) The EMBL-EBI Job Dispatcher sequence analysis tools framework in 2024. Nucleic Acids Res., 52, W521–W525.

104. Waterhouse, A.M., Procter, J.B., Martin, D.M.A., Clamp, M. and Barton, G.J. (2009) Jalview Version 2-A multiple sequence alignment editor and analysis workbench. Bioinformatics, 10.1093/bioinformatics/btp033.

