## Supplementary Tables and Figures for "Structural mechanism of mRNA decoding by mammalian GTPase GTPBP1"

| | Models | | | | | | | | | | Supplementary maps | | |
|---|---|---|---|---|---|---|---|---|---|---|---|---|---|
| | Structure III | Structure IIa | Structure IIb | Structure IIc | Structure IId | Structure IV | Structure Ia | Structure Ib | Structure Ic | Structure V | Structure IV-like<br>5 min GTP-dataset | Structure III<br>composite mask | Structure IV<br>composite mask |
| **Cryo-EM data** | | | | | | | | | | | | | |
| Voltage (kV) | 300 | 300 | 300 | 300 | 300 | 300 | 300 | 300 | 300 | 300 | 300 | 300 | 300 |
| Electron exposure (e–/Å$^2$) | 29.8 | 29.8 | 29.8 | 29.8 | 29.8 | 29.8 | 30 | 30 | 30 | 30 | 30 | 29.8 | 29.8 |
| Defocus range (μm) | 0.5-1.5 | 0.5-1.5 | 0.5-1.5 | 0.5-1.5 | 0.5-1.5 | 0.5-1.5 | 0.5-1.5 | 0.5-1.5 | 0.5-1.5 | 0.5-1.5 | 0.5-1.5 | 0.5-1.5 | 0.5-1.5 |
| Pixel size (Å) | 1.162 | 1.162 | 1.162 | 1.162 | 1.162 | 1.167 | 1.162 | 1.162 | 1.162 | 1.162 | 1.162 | 1.162 | 1.167 |
| Symmetry imposed | C1 | C1 | C1 | C1 | C1 | C1 | C1 | C1 | C1 | C1 | C1 | C1 | C1 |
| Initial particle images (no.) | 318381 | 318381 | 318381 | 318381 | 318381 | 1029824 | 519352 | 519352 | 519352 | 519352 | 519352 | 318381 | 1029824 |
| Final particle images (no.) | 8990 | 8005 | 7906 | 7386 | 7140 | 9969 | 10400 | 10267 | 11317 | 15402 | 5287 | 8990 | 9969 |
| Map resolution (Å) | 3.0 | 3.0 | 3.0 | 3.1 | 3.0 | 2.9 | 2.9 | 2.9 | 2.9 | 3.0 | 3.0 | 3.7 | 4.1 |
| Map sharpening B factor (Å$^2$) | – | – | – | – | – | – | – | – | – | – | – | -50 | -50 |
| **Composition** | | | | | | | | | | | | | |
| Chains | 85 | 85 | 85 | 85 | 85 | 85 | 83 | 83 | 83 | 84 | | | |
| Atoms | 219622<br>(Hydrogens: 0) | 219611<br>(Hydrogens: 0) | 219611<br>(Hydrogens: 0) | 219611<br>(Hydrogens: 0) | 219611<br>(Hydrogens: 0) | 219607<br>(Hydrogens: 0) | 213971<br>(Hydrogens: 0) | 213971<br>(Hydrogens: 0) | 213971<br>(Hydrogens: 0) | 215594<br>(Hydrogens: 0) | | | |
| Residues | Protein: 11835<br>Nucleotide: 5802 | Protein: 11835<br>Nucleotide: 5802 | Protein: 11835<br>Nucleotide: 5802 | Protein: 11835<br>Nucleotide: 5802 | Protein: 11835<br>Nucleotide: 5802 | Protein: 11835<br>Nucleotide: 5802 | Protein: 11323<br>Nucleotide: 5727 | Protein: 11323<br>Nucleotide: 5727 | Protein: 11323<br>Nucleotide: 5727 | Protein: 11322<br>Nucleotide: 5802 | | | |
| Water | 0 | 0 | 0 | 0 | 0 | 0 | 0 | 0 | 0 | 0 | | | |
| Ligands | ZN: 6<br>K: 1<br>GCP: 1<br>MG: 1<br>SPD: 1 | ZN: 6<br>K: 1<br>GCP: 1<br>MG: 1<br>SPD: 1 | ZN: 6<br>K: 1<br>GCP: 1<br>MG: 1<br>SPD: 1 | ZN: 6<br>K: 1<br>GCP: 1<br>MG: 1<br>SPD: 1 | ZN: 6<br>K: 1<br>GCP: 1<br>MG: 1<br>SPD: 1 | ZN: 6<br>K: 1<br>GDP: 1<br>MG: 1<br>SPD: 1 | ZN: 6<br>SPD: 1 | ZN: 6<br>SPD: 1 | ZN: 6<br>SPD: 1 | ZN: 6<br>SPD: 1 | | | |
| **Bonds (RMSD)** | | | | | | | | | | | | | |
| Length (Å) (# > 4σ) | 0.007 (0) | 0.008 (0) | 0.008 (0) | 0.008 (0) | 0.007 (0) | 0.007 (0) | 0.007 (0) | 0.008 (0) | 0.008 (0) | 0.008 (0) | | | |
| Angles (Å) (# > 4σ) | 0.995 (138) | 1.003 (141) | 1.002 (157) | 1.011 (141) | 0.978 (129) | 0.992 (125) | 1.006 (145) | 1.018 (154) | 1.013 (125) | 1.017 (133) | | | |
| **MolProbity score** | 1.38 | 1.36 | 1.43 | 1.42 | 1.41 | 1.37 | 1.36 | 1.38 | 1.37 | 1.4 | | | |
| **Clash score** | 4.64 | 4.59 | 4.82 | 4.87 | 4.87 | 4.47 | 4.66 | 4.52 | 4.66 | 4.94 | | | |
| **Ramachandran plot (%)** | | | | | | | | | | | | | |
| Outliers | 0 | 0 | 0.01 | 0 | 0 | 0.01 | 0 | 0 | 0 | 0 | | | |
| Allowed | 2.79 | 2.67 | 3.02 | 2.97 | 2.87 | 2.79 | 2.66 | 2.83 | 2.67 | 2.75 | | | |
| Favored | 97.21 | 97.33 | 96.97 | 97.03 | 97.13 | 97.2 | 97.34 | 97.17 | 97.33 | 97.25 | | | |
| **Rama-Z (Ramachandran plot Z-score, RMSD)** | | | | | | | | | | | | | |
| whole (N = 11660) | -1.02 (0.07) | -0.90 (0.07) | -0.91 (0.07) | -0.92 (0.07) | -0.91 (0.07) | -0.85 (0.07) | -0.77 (0.07) | -0.88 (0.07) | -0.89 (0.07) | -0.89 (0.07) | | | |
| helix (N = 3957) | -1.16 (0.07) | -1.06 (0.07) | -1.04 (0.07) | -1.10 (0.07) | -1.12 (0.07) | -1.11 (0.07) | -0.97 (0.07) | -1.06 (0.07) | -1.04 (0.07) | -1.07 (0.07) | | | |
| sheet (N = 1935) | -0.05 (0.11) | 0.04 (0.11) | -0.09 (0.11) | -0.04 (0.11) | 0.06 (0.11) | 0.09 (0.11) | 0.05 (0.12) | -0.16 (0.11) | -0.20 (0.11) | -0.12 (0.12) | | | |
| loop (N = 5768) | -0.38 (0.08) | -0.31 (0.08) | -0.29 (0.08) | -0.30 (0.08) | -0.31 (0.08) | -0.24 (0.08) | -0.19 (0.08) | -0.20 (0.08) | -0.22 (0.08) | -0.22 (0.08) | | | |
| **Rotamer outliers (%)** | 0 | 0.02 | 0.03 | 0.04 | 0.05 | 0.06 | 0.03 | 0.11 | 0.01 | 0.02 | | | |
| **Cβ outliers (%)** | NA | NA | NA | NA | NA | NA | NA | NA | NA | NA | | | |
| **Peptide plane (%)** | | | | | | | | | | | | | |
| Cis proline/general | 2.9/0.0 | 2.9/0.0 | 2.9/0.0 | 2.9/0.0 | 2.9/0.0 | 2.9/0.0 | 2.8/0.0 | 2.8/0.0 | 2.8/0.0 | 2.8/0.0 | | | |
| Twisted proline/general | 0.0/0.0 | 0.0/0.0 | 0.0/0.0 | 0.0/0.0 | 0.0/0.0 | 0.0/0.0 | 0.0/0.0 | 0.0/0.0 | 0.0/0.0 | 0.0/0.0 | | | |
| **CaBLAM outliers (%)** | 1.32 | 1.25 | 1.3 | 1.38 | 1.23 | 1.38 | 1.21 | 1.36 | 1.39 | 1.33 | | | |
| **ADP (B-factors)** | | | | | | | | | | | | | |
| Iso/Aniso (#) | 219622/0 | 219611/0 | 219611/0 | 219611/0 | 219611/0 | 219607/0 | 213971/0 | 213971/0 | 213971/0 | 215594/0 | | | |
| min/max/mean | | | | | | | | | | | | | |
| Protein | 7.89/235.97/72.86 | 8.29/321.46/74.26 | 19.24/261.32/75.10 | 11.23/303.40/75.80 | 14.63/303.67/73.16 | 16.60/252.09/69.48 | 14.57/241.22/64.56 | 11.20/240.80/64.01 | 7.72/236.59/65.09 | 13.25/226.48/67.73 | | | |
| Nucleotide | 0.00/310.41/77.54 | 0.31/336.51/75.66 | 0.44/344.19/77.62 | 0.41/370.80/82.10 | 0.00/347.16/76.28 | 0.00/331.09/72.43 | 0.00/319.99/71.88 | 0.50/302.78/69.96 | 0.00/308.46/71.14 | 0.05/311.32/74.25 | | | |
| Ligand | 24.50/210.55/110.31 | 24.25/196.21/109.74 | 23.99/213.35/108.51 | 30.36/217.67/132.89 | 24.90/208.32/103.28 | 18.96/203.82/122.72 | 21.70/179.38/122.75 | 21.33/172.52/96.84 | 23.11/174.94/117.17 | 25.20/263.27/135.86 | | | |
| Water | --- | --- | --- | --- | --- | --- | --- | --- | --- | --- | | | |
| **Occupancy** | | | | | | | | | | | | | |
| Mean | 1 | 1 | 1 | 1 | 1 | 1 | 1 | 1 | 1 | 1 | | | |
| occ = 1 (%) | 99.97 | 99.98 | 99.98 | 99.98 | 99.98 | 99.98 | 100 | 100 | 100 | 99.98 | | | |
| 0 < occ < 1 (%) | 0.02 | 0.01 | 0.01 | 0.01 | 0.01 | 0.01 | 0 | 0 | 0 | 0.01 | | | |
| occ > 1 (%) | 0 | 0 | 0 | 0 | 0 | 0 | 0 | 0 | 0 | 0 | | | |
| **Model vs. Data** | | | | | | | | | | | | | |
| CC (mask) | 0.88 | 0.87 | 0.87 | 0.87 | 0.86 | 0.88 | 0.88 | 0.88 | 0.88 | 0.87 | | | |
| CC (box) | 0.81 | 0.81 | 0.81 | 0.81 | 0.8 | 0.81 | 0.8 | 0.8 | 0.81 | 0.8 | | | |
| CC (peaks) | 0.79 | 0.79 | 0.79 | 0.79 | 0.78 | 0.8 | 0.8 | 0.8 | 0.8 | 0.79 | | | |
| CC (volume) | 0.87 | 0.87 | 0.87 | 0.86 | 0.86 | 0.87 | 0.87 | 0.87 | 0.88 | 0.87 | | | |
| Mean CC for ligands | 0.83 | 0.8 | 0.8 | 0.79 | 0.74 | 0.76 | 0.84 | 0.83 | 0.83 | 0.83 | | | |
| **RNA** | | | | | | | | | | | | | |
| good pucker (%) | 95.5 | 95.8 | 95.8 | 95.6 | 95.4 | 95.5 | 95.4 | 95.6 | 95.4 | 95.5 | | | |
| good backbone (%) | 85.3 | 85.7 | 85.4 | 85.1 | 85.2 | 85.4 | 85.6 | 85.4 | 85.2 | 84.8 | | | |

**Supplementary Table 1**. Cryo-EM data and structure statistics.

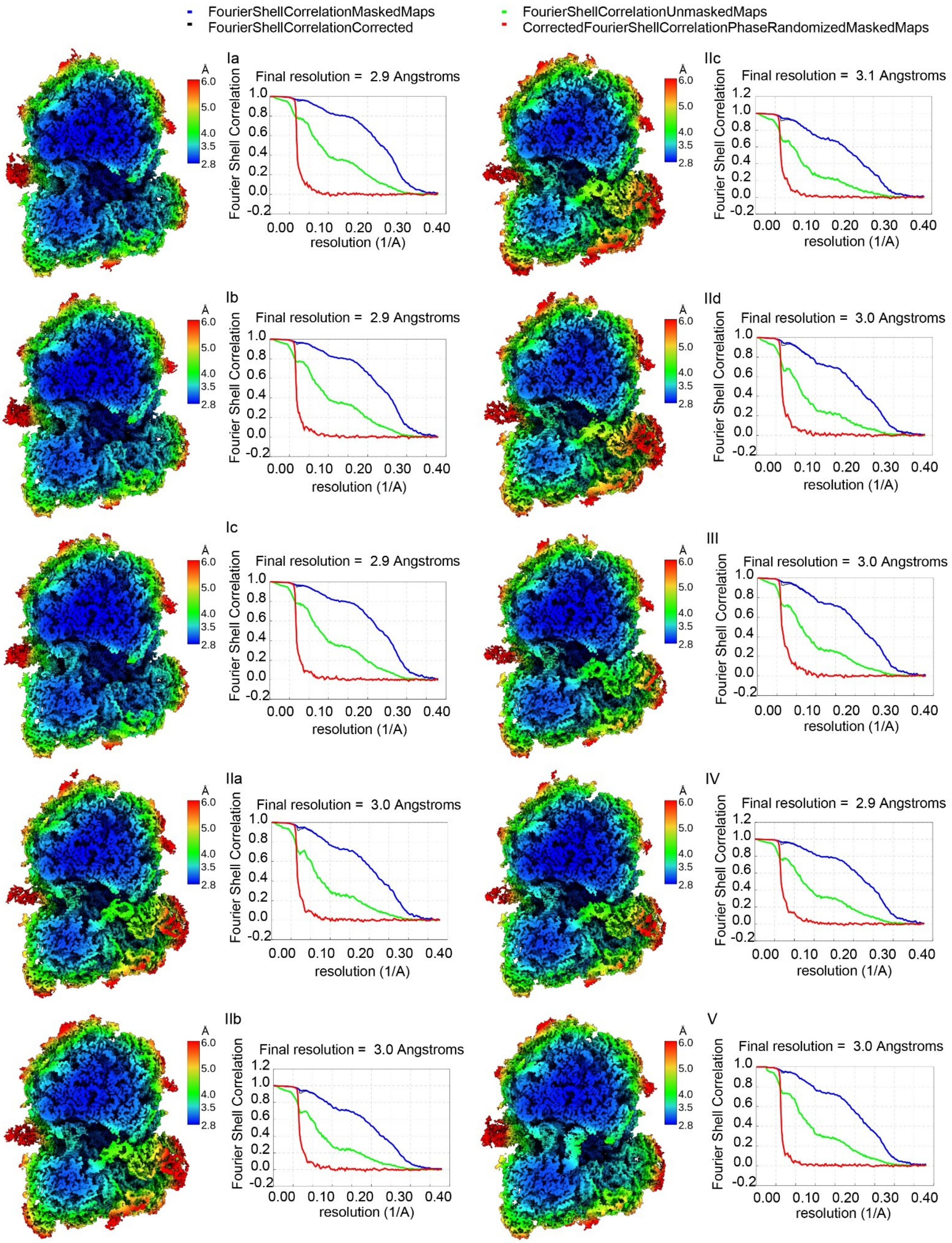


**Supplementary Figure 1**. Local resolutions and Fourier shell correlation curves for cryo-EM maps Ia through V.

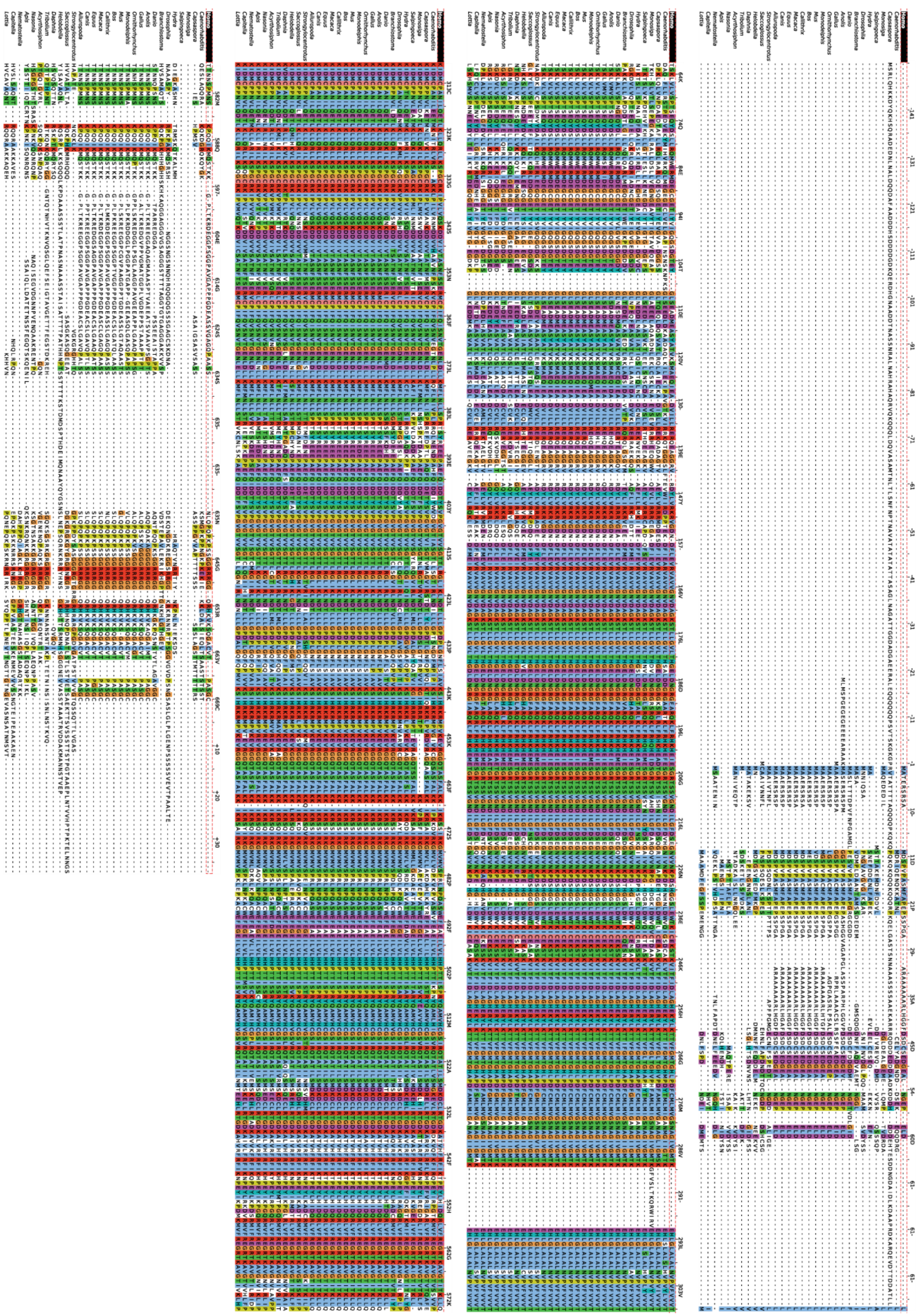

**Supplementary Figure 2a**. GTPBP1 protein sequence alignment.

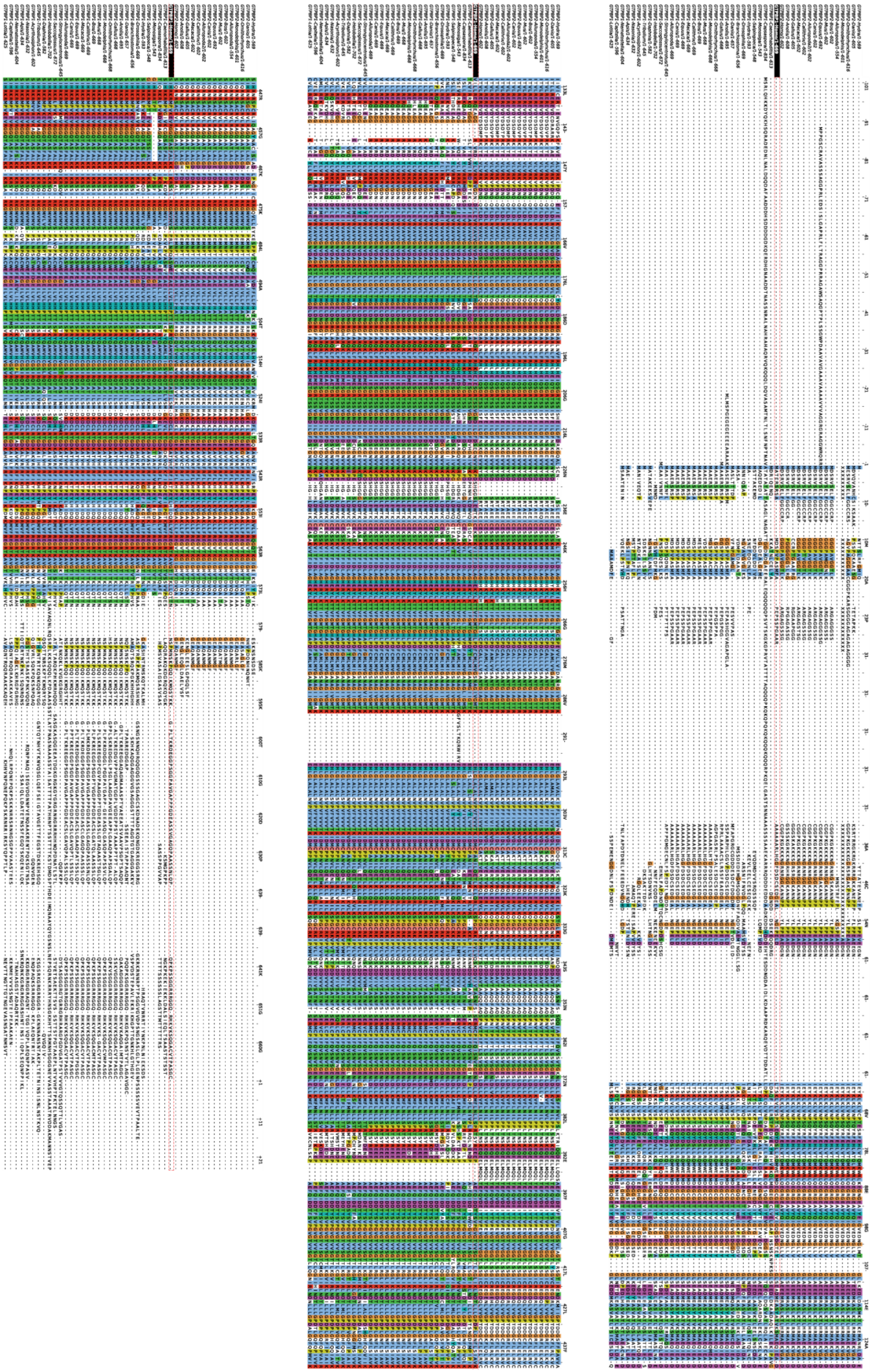

**Supplementary Figure 2b**. GTPBP1 and GTPBP2 protein sequence alignment.

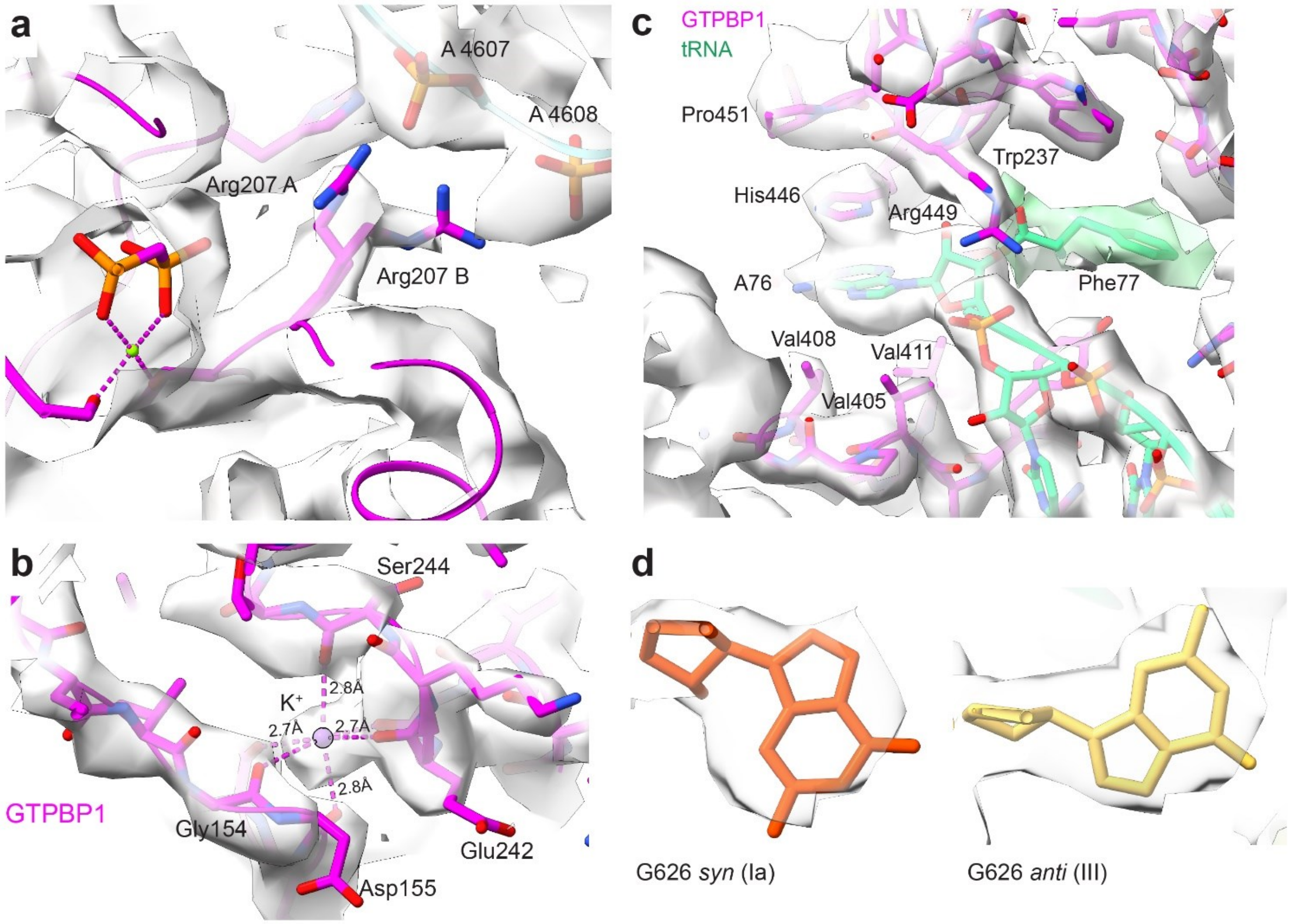


**Supplementary Figure 3**. Examples of local cryo-EM densities. **a**) Two alternative rotamers of Arg207 in the GTPase center of GTPBP1 (Structure III). **b**) $K^+$-binding pocket of GTPBP1 (Structure III, from the stack refined using a composite mask). **c**) tRNA-binding pocket of GTPBP1 (Structure III). The density around Phe77 is shown at a lower contour level (0.01) than the rest of the map (0.016). **d**) G626 in the *syn* and *anti* conformations in Structures Ia and III, respectively.

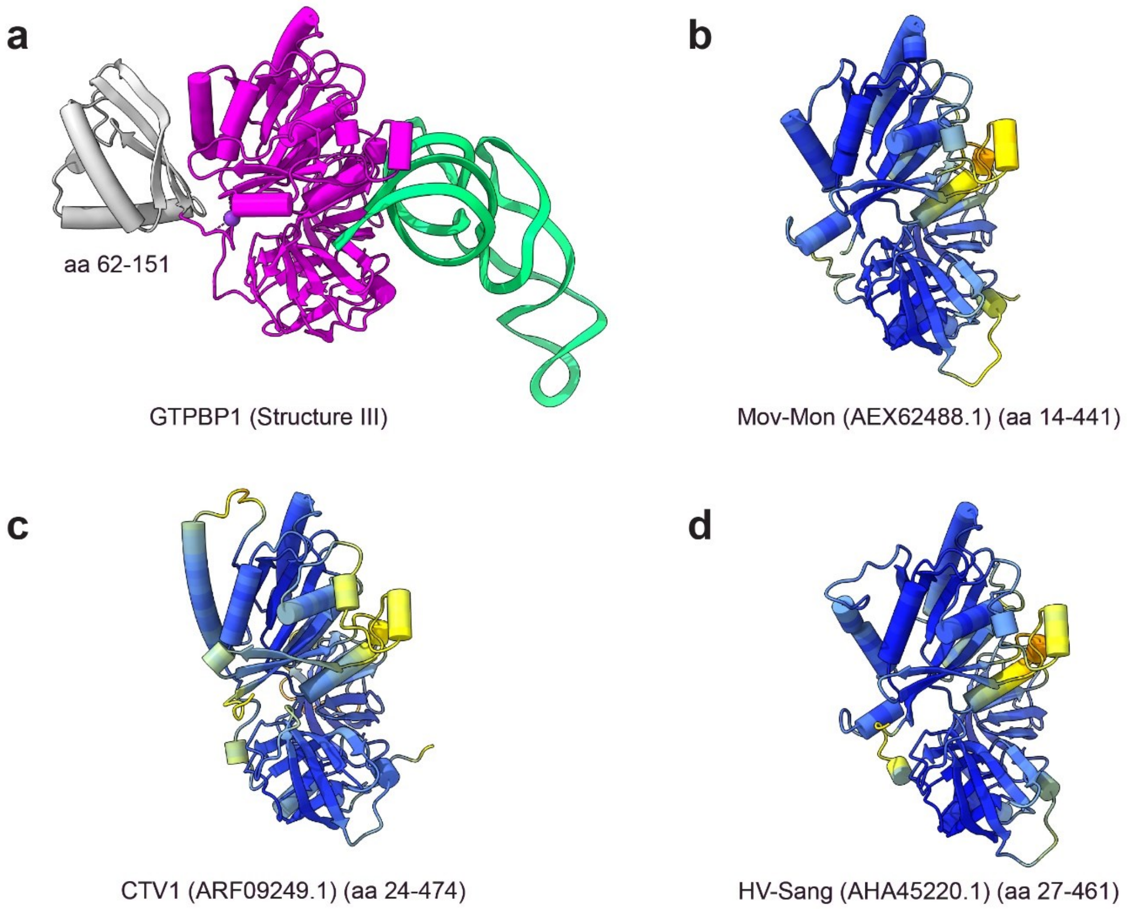


**Supplementary Figure 4**. Comparison of human GTPBP1 with viral GTPBP1-like proteins. **a**) GTPBP1•aa-tRNA (Structure III) with; amino acids deleted in Zinoviev *et al* (1) shown in gray. **b-d**) Alphafold (2) models of different viral GTPBP1-like proteins, colored by per-residue model confidence score (pLDDT) from red (low) to blue (high).

1. Zinoviev,A., Kuroha,K., Pestova,T. V. and Hellen,C.U.T. (2019) Two classes of EF1-family translational GTPases encoded by giant viruses. *Nucleic Acids Res.*, 10.1093/nar/gkz296.
2. Jumper,J., Evans,R., Pritzel,A., Green,T., Figurnov,M., Ronneberger,O., Tunyasuvunakool,K., Bates,R., Žídek,A., Potapenko,A., *et al.* (2021) Highly accurate protein structure prediction with AlphaFold. *Nature*, 10.1038/s41586-021-03819-2.

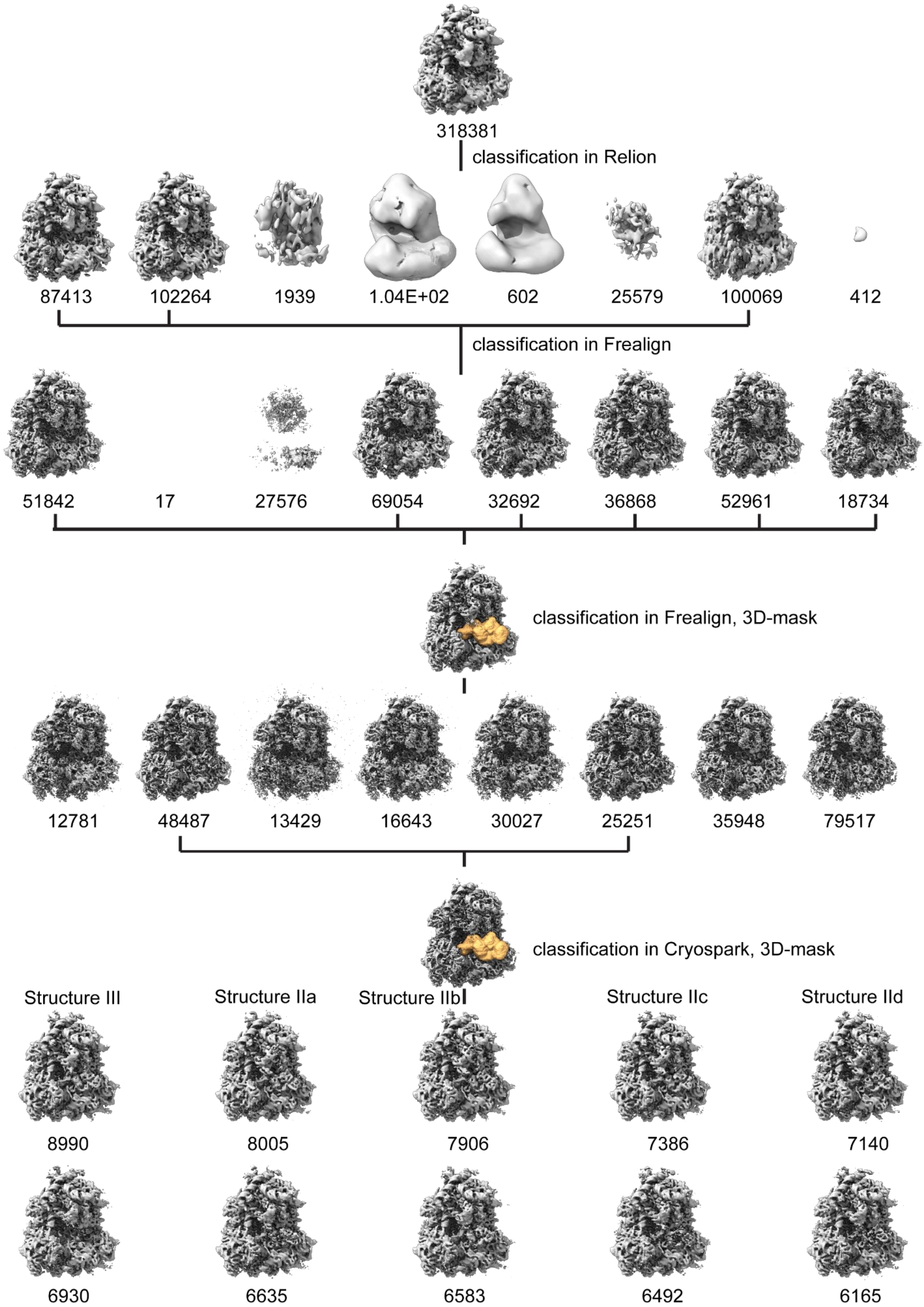


**Supplementary Figure 5a**. Scheme of the classification of the GDPCP dataset. Number of particles is shown for each reconstruction; 3D masks are shown as yellow volumes.

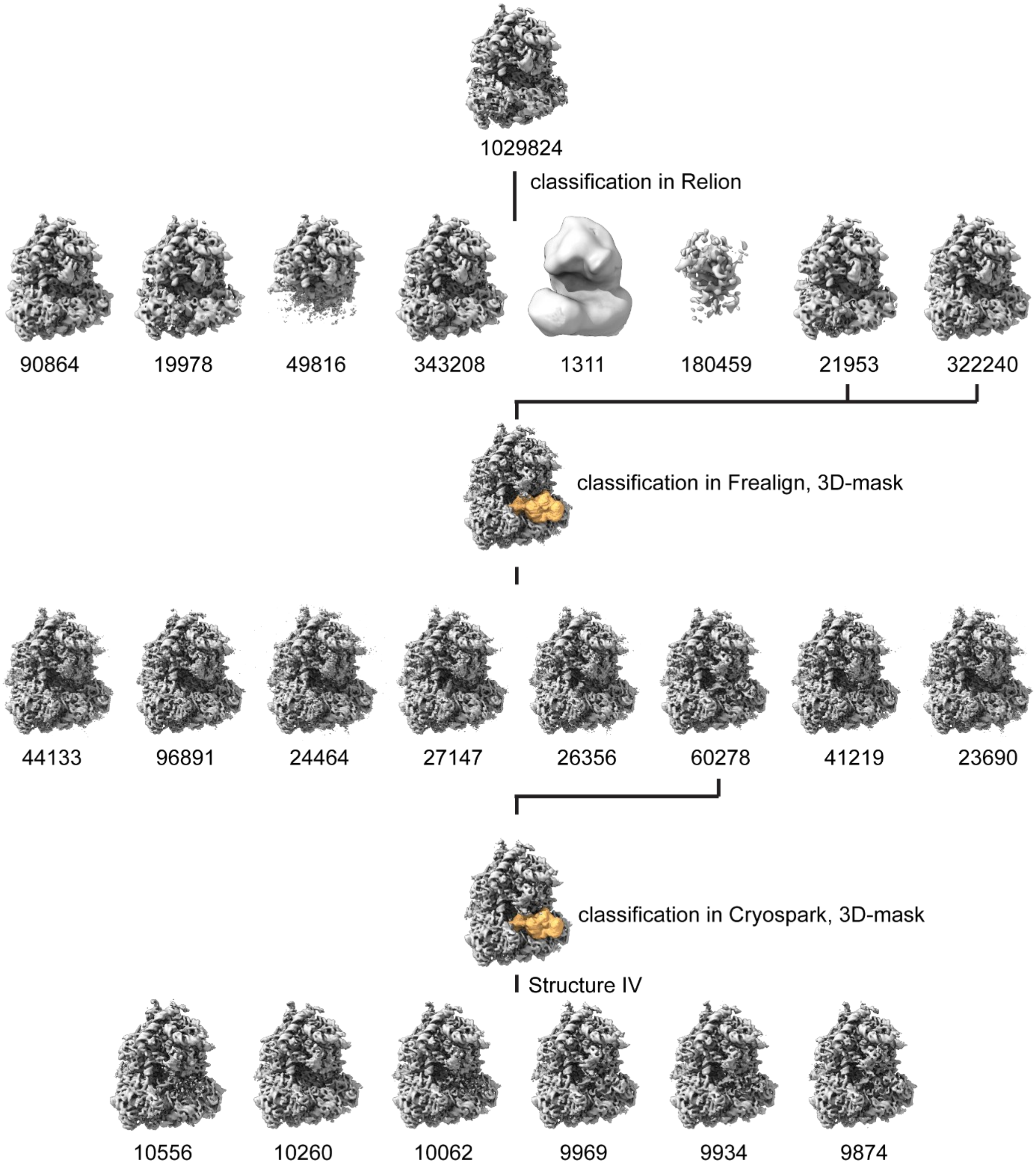


**Supplementary Figure 5b**. Scheme of the classification of the GTP 1-minute dataset. Number of particles is shown for each reconstruction; 3D masks are shown as yellow volumes.

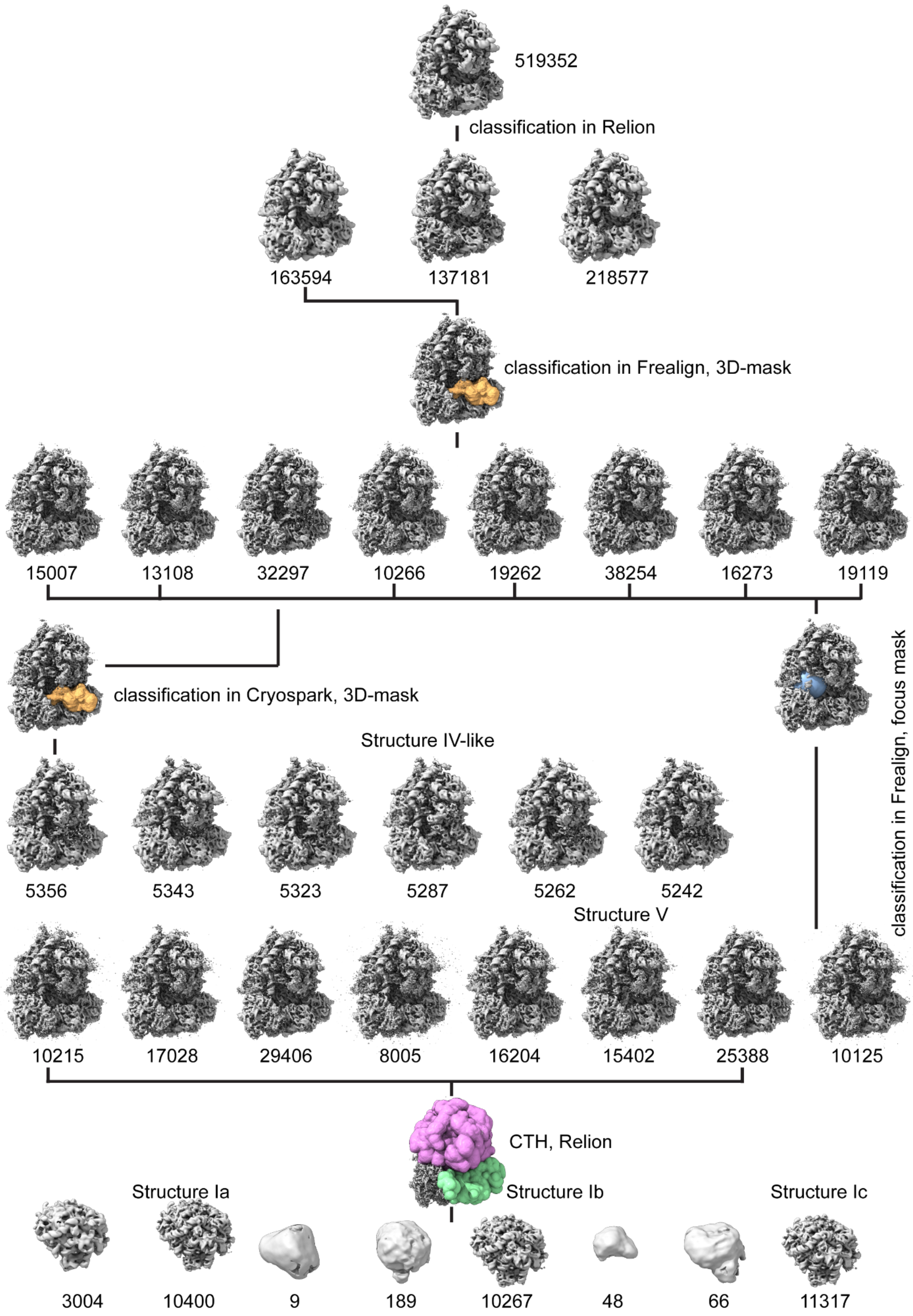


**Supplementary Figure 5c**. Scheme of the classification of the GTP 5-minute dataset. Number of particles is shown for each reconstruction; 3D masks are shown as yellow, green and pink volumes.

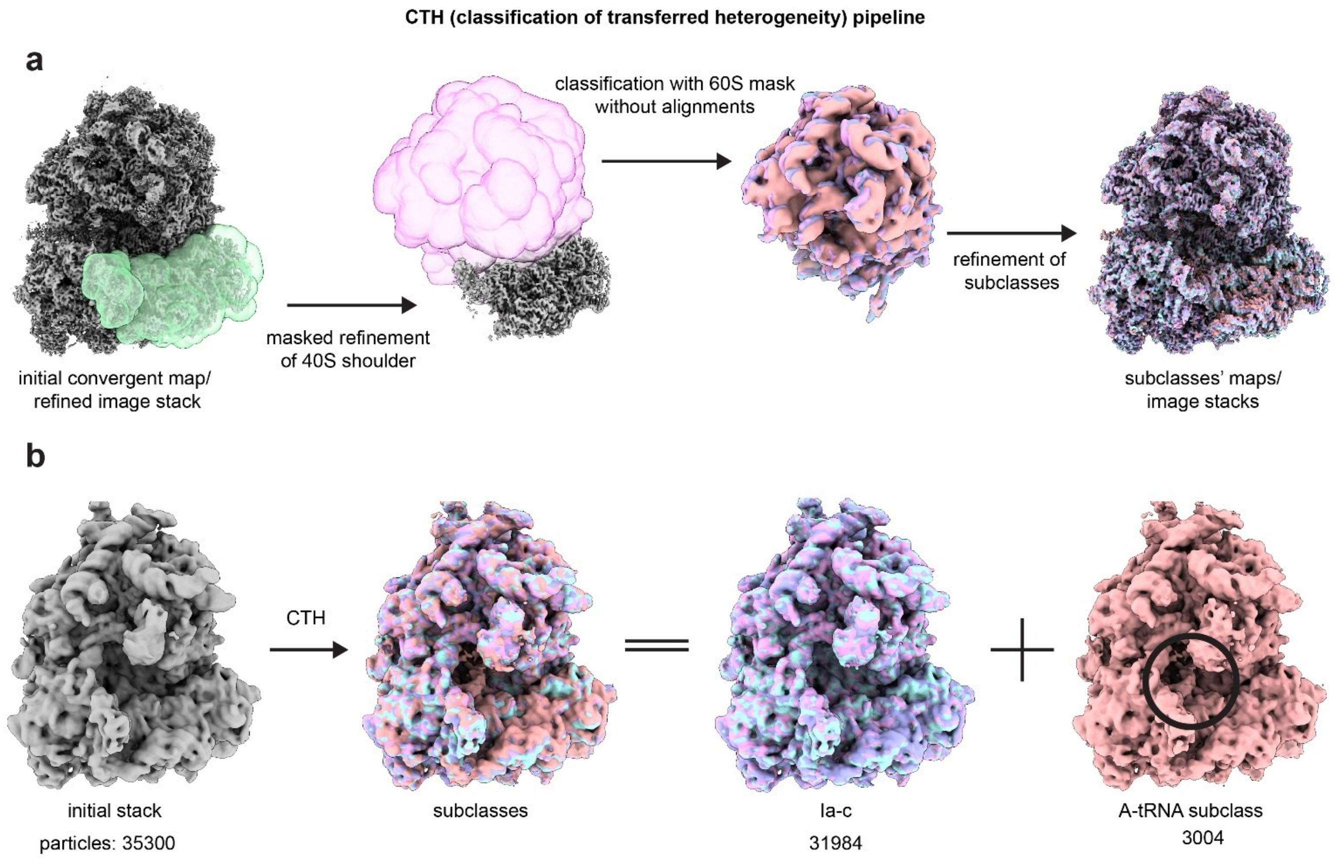


**Supplementary Figure 6**. The CTH (classification of transferred heterogeneity) approach for improved classification of local cryo-EM densities. **a)** The CTH workflow. 3D masks are shown as green and pink volumes. **b)** CTH allows for the detection of low-abundance A-tRNA subclasses (A-tRNA is highlighted by the black circle).
